# Transcriptomic response in pyroxsulam-resistant and susceptible *Bromus sterilis* identified three distinct mechanisms of resistance

**DOI:** 10.1101/2023.07.14.548957

**Authors:** Madhab Kumar Sen, Katerina Hamouzová, Nawaporn Onkokesung, Julio Menendez, Joel Torra, Pavlina Košnarová, Gothandapani Sellamuthu, Aayushi Gupta, Rohit Bharati, Vishma Pratap Sur, Amit Roy, Josef Soukup

## Abstract

*Bromus sterilis* has evolved into a more predominant weed in the Czech Republic’s winter wheat fields, owing largely to the widespread application of pyroxsulam for its management. In this study, we report a biotype that has developed resistance to pyroxsulam and has also shown cross-resistance to other herbicides. Although no differences in ploidy levels or no mutations of acetolactate synthase (ALS) were detected, a significant elevation of ALS enzyme activity was observed in the R biotype. Through combined analysis of enzyme inhibition and total transcript expression (RNA-Seq), we have identified differentially expressed transcripts that potentially contribute to pyroxsulam metabolism. Furthermore, we observed a significant increase in the expression of genes involved in redox mechanisms and transporters that could contribute to enhanced resistance to pyroxulam in the R biotype. Our results present a novel understanding of herbicide resistance in *B. sterilis* through three distinct resistance mechanisms (*ALS* gene overexpression, enhanced metabolism and reduced translocation) without mutation in the herbicide target protein. This understanding is the foundation for improving management strategies for herbicide resistant *B. sterilis*.

## 1. Introduction

Weeds pose a significant threat to agricultural production systems as they compete with crops for resources such as nutrients, sunlight, and water, resulting in yield losses (Chauhan, 2020; Flessner *et al*., 2021). An application of synthetic chemical herbicides is a common and cost- effectuve approach to manage agriculture weeds. However, the repated application of herbicides lead to the development of herbicide resistance. In fact, a global survey has identified 268 species (154 dicots and 114 monocots) that have developed resistance against 165 different herbicides (https://www.weedscience.org/Home.aspx). The acetolactate synthase (ALS)- inhibiting herbicide class is one of the herbicides that have been widely used gloablly. These herbicides offer broad-spectrum weed control by targeting the ALS enzyme in weeds. ALS is an essential enzyme involved in the biosynthesis of branched-chain amino acids (Sen *et al*., 2021). Among 58 ALS herbicides based on Herbicide Resistance Action Committee classification, pyroxulam has been commonly used to control grass weeds in Europe, North America and Australia. Despite the initial success of pyroxsulam in effective weed management, the overuse has led to the evolution of herbicide resistance in major weed species. These include *Amaranthus* spp. (Wang *et al*., 2022), *Lolium multiflorum* L. (Kaya Altop *et al*., 2022) and *Bromus sterilis* L. (barren brome) (Sen *et al*., 2021).

*Bromus sterilis*, an annual or biennial grass weed, is well-known for its detrimental effects on winter wheat productivity, causing yield reductions of up to 60% (Sen *et al*., 2021). Moreover, *B. sterilis* can also disrupt harvesting processes and even lead to lodging issues (Moray et al., 2003). In the Czech Republic (CZ), this troublesome weed is prevalent in lowland areas, and its management primarily relies on ALS-inhibiting herbicides such as pyroxsulam and propoxycarbazone (Sen *et al*., 2021). Apart from CZ, *B. sterilis* herbicide resistant biotypes have also been reported in France (Heap, 2023), Germany (Heap, 2023), and the United Kingdom (Davis *et al*., 2020). Typically, herbicide resistance in weeds can evolve through two primary mechanisms: target site resistance (TSR) and non-target site resistance (NTSR) (Gaines *et al*., 2020). TSR mechanisms occur when genetic changes in the weed modify the herbicide’s target site through point mutations or increase target gene copy numbers, while NTSR mechanisms involve the mechanisms that prevent the herbicidal molecules from reaching their target site (Gaines *et al*., 2020). Based on current understanding, NTSR mechanisms include increased herbicide detoxification, reduced penetration, and absorption. These mechanisms are associated with complex gene families such as cytochrome P450s (Cyp450), glutathione S-transferases (GSTs), glycosyl transferases (GTs), and ATP-binding cassette (ABC) transporters (Torra & Alcántara-de la Cruz, 2022; Délye, 2013). While many isoforms of these gene families have been identified as being linked with herbicide resistance, there are likely numerous others yet to be discovered (Dimaano & Iwakami, 2021). Nevertheless, evolution of the herbicide resistance in weeds by accumulating several mechanisms serves as a notable example of plants adapting against human selective pressures. It is crucial to understand these mechanisms in *B. sterilis* to mitigate the consequences of herbicide resistance on crop production.

To date, *ALS* gene overexpression and NTSR mechanisms (via Cyp450s and GSTs) have been reported to be the underlying resistance mechanisms in *B. sterilis* (Sen *et al*., 2021; Davis *et al*., 2020). However, conclusive studies to elucidate resistance mechanisms are still lacking. In this study, we performed RNA-seq analysis on a pyroxsulam-resistant biotype of *B. sterilis* collected from a winter wheat field in CZ. The differential expression analysis suggests the involvement of three different mechanisms in conferring resistance to pyroxsulam in the R biotype. We also investigated cross-resistance to other herbicides. Based on our findings, we strongly recommend an immediate cessation of pyroxsulam use and the adoption of alternative herbicides within an integrated weed management approach to effectively manage this species.

## 2. Materials and methods

### 2.1 Plant materials and herbicide-response experiments

Two biotypes, namely the susceptible (S) and the resistant (R) biotypes, were used in this study. The S biotype was collected from a winter wheat field in the Jihomoravský (Vyškov) region (49.1219997N, 16.8183625E), while the resistant (R) biotype was collected from a winter wheat field in the Ústecký (Louny) region of CZ (50.2784172N, 13.5348256E). Seeds were collected in bulk from a minimum of 100 plants and stored in darkness at room temperature until further use. Approximately 10 seeds were sown in pots (∼343Lcm^3^), filled with chernozem soil [clay content 46% (loamy soil), soil pH (potassium chloride) 7.5, sorption capacity of soil: 209Lmmol (+), 87LmgLkg^−1^ phosphorus, 203LmgLkg^−1^ potassium, 197LmgLkg^−1^ magnesium, 8073LmgLkg^−1^ calcium]. The pot experiments were conducted in an open-air vegetation hall with a roof-top, and the seedlings were watered and fertilized as required. The herbicides were treated at a two-three leaves stage, with a laboratory spray chamber supplied with a Lurmark 015F80 nozzle with spray volume of 250LLLha^−1^ and pressure 120LkPa. The list of doses of the herbicides are given in table 1. Pyroxsulam was applied at the rates of 0.05925, 0.1875, 0.5925, 1.875, 5.925, 18.75, 59.25, 187.5 and 592.5 g a.i. (active ingredient) ha^−1^. All the pot experiments were done with four-pot replicates for each rate. Malathion (Malathion, PESTANAL®, analytical standard, Sigma-Aldrich, Merck Group, St Louis, MO, USA) was applied at 1000LgLha^−1^ of active ingredient, while 4-chloro-7-nitrobenzoxadiazole (NBD-Cl, 97%, Sigma-Aldrich, Merck Group, St Louis, MO, USA) was applied at 270LgLha^−1^ of active ingredient. Malathion and NBD-Cl were applied at 1h and 2d prior to herbicide treatment, respectively [3, 13]. The dose-response curves and the GR50 (50% growth inhibition) were calculated using the R-Studio program (https://www.r-project.org/). The dose–response curves were plotted using a non–linear regression model, and the resistance factor (RF) ratios were determined based on the GR50 values. The cross-resistance results were verified using ANOVA in R-Studio program, and Tukey’s post hoc analysis was employed for pairwise comparisons at a significance level of 5%.

**Table 1:**
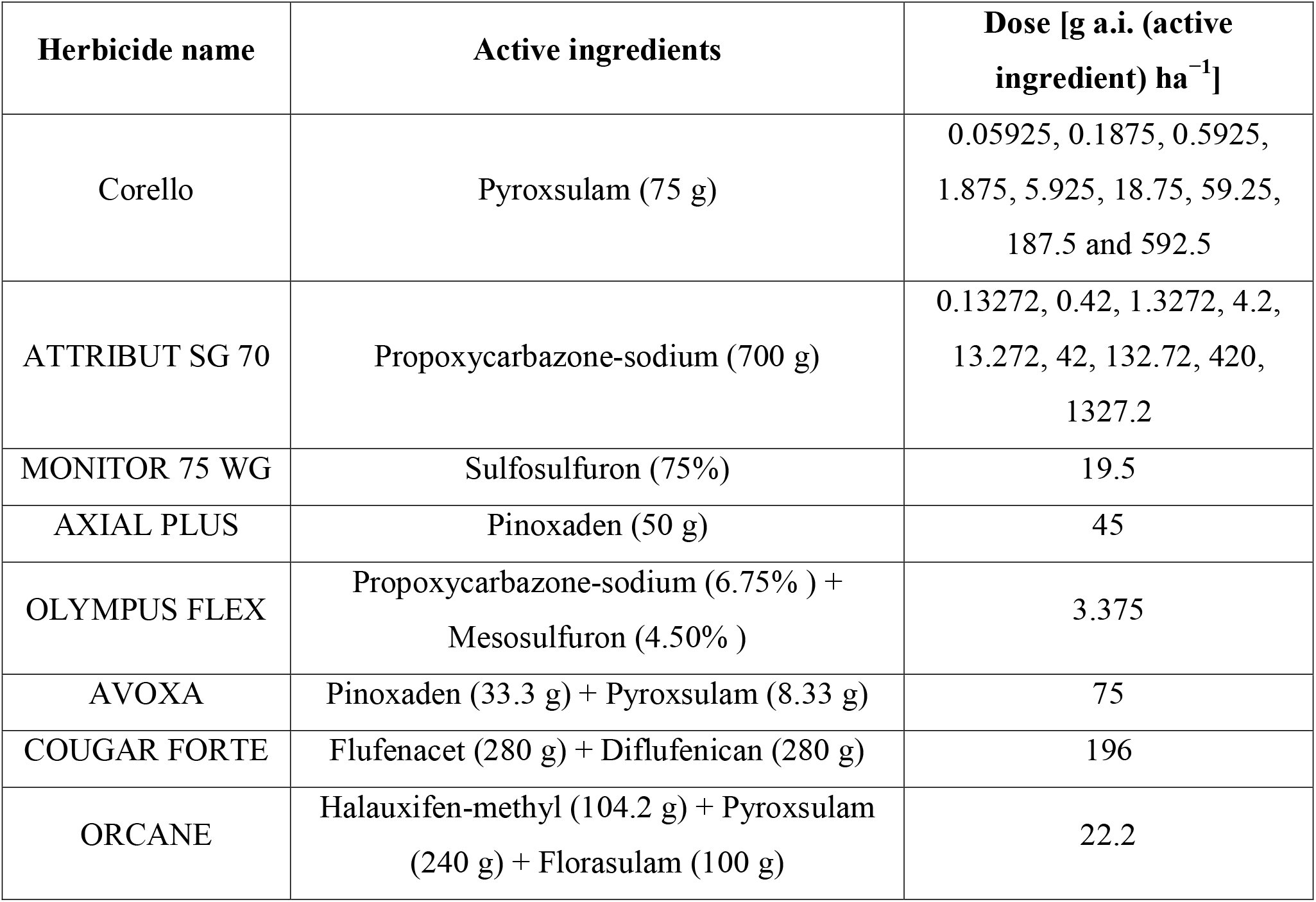
List of doses used in the study.

| Herbicide name | Active ingredients | Dose [g a.i. (active ingredient) ha<sup>-1</sup>] |
| --- | --- | --- |
| Corello | Pyroxsulam (75 g) | 0.05925, 0.1875, 0.5925, 1.875, 5.925, 18.75, 59.25, 187.5 and 592.5 |
| ATTRIBUT SG 70 | Propoxycarbazone-sodium (700 g) | 0.13272, 0.42, 1.3272, 4.2, 13.272, 42, 132.72, 420, 1327.2 |
| MONITOR 75 WG | Sulfosulfuron (75%) | 19.5 |
| AXIAL PLUS | Pinoxaden (50 g) | 45 |
| OLYMPUS FLEX | Propoxycarbazone-sodium (6.75% ) + Mesosulfuron (4.50% ) | 3.375 |
| AVOXA | Pinoxaden (33.3 g) + Pyroxsulam (8.33 g) | 75 |
| COUGAR FORTE | Flufenacet (280 g) + Diflufenican (280 g) | 196 |
| ORCANE | Halauxifen-methyl (104.2 g) + Pyroxsulam (240 g) + Florasulam (100 g) | 22.2 |

### 2.2 Flow cytometry analysis

The flow cytometry procedure followed the method described by Bharati et al. (2023) with slight modifications (Bharati *et al*., 2023). Briefly, the leaf segments of approximately 0.5 cm^2^ from both R and S biotypes were placed in a Petri dish and finely chopped using a razor blade in 1 mL of Otto I buffer (0.1 M citric acid monohydrate + 0.5% Tween 20). Subsequently, the suspension was filtered through a 50 µm nylon mesh and the filtrate obtained was mixed with 1 mL of Otto II buffer (0.4 M Na_2_HPO_4_·12 H_2_O) containing 2 μg/mL of DAPI, a fluorescent dye. Finally, the resulting solutions were analyzed using a Partec PAS flow cytometer (Partec GmbH, Münster, Germany) equipped with a high-pressure mercury arc lamp, and histograms displaying the relative DNA content were generated using the Flomax software package (Version 2.3).

### 2.3 ALS enzyme activity

ALS activity was determined by measuring the production of acetolactate, which is formed after decarboxylation into acetoin in the presence of acid, as described in Osuna et al., 2002. For the assay, two grams of young leaf tissue were collected from each population (R and S). Plant tissue used for the assays was obtained from the S and R plants, which were grown under the same conditions as described in section 2.1. The ALS enzyme activity assay followed the method outlined by Palma-Bautista *et al*., 2023, and the I50 (the concentration required to reduce ALS activity by 50%) was calculated according to Cruz-Hipolito *et al*., 2009. To evaluate ALS enzyme activity, enzyme extract was added to a fresh assay buffer consisting of 1M KH_2_PO_4_/K_2_HPO_4_, pH 7.5; 0.5M sodium pyruvate; 0.1M MgCl_2_, 50 mM TPP; and 1 mm FAD, along with increasing concentrations of herbicides. The assay utilized a technical grade of pyroxsulam at various concentrations ranging from 10^−2^ to 10^8^ nm of active ingredient (Hamouzová et al., 2011).

### 2.4 Gene sequencing, overexpression and copy number variation studies

Shock-frozen leaf tissues weighing approximately 80 mg per sample from R and S plants were used for the extraction of genomic DNA (gDNA) and total RNA. The DNeasy Plant Mini Kit (QIAGEN, Hilden, Germany) was employed for gDNA extraction, following the manufacturer’s instructions. RNA extraction was carried out using the RNeasy Mini Kit (QIAGEN, Hilden, Germany), and cDNA synthesis was performed with the High Capacity cDNA Reverse Transcription Kit (Applied Biosystems, Foster City, CA, USA). Approximately 1 µg of RNA template was used for cDNA synthesis. PCR was conducted using a C1000 thermocycler (Bio-Rad, Hercules, CA, USA), utilizing 50 ng of total gDNA per reaction. The relative copy number and expression experiments were performed using StepOne™ Real-Time PCR System (Applied Biosystems) with genomic DNA (∼10 ng) and complementary DNA (cDNA, ∼10 ng) as templates. Detailed information regarding the primers and PCR and qRT-PCR procedures can be found in Sen et al., 2021. The relative ALS copy number and expression values were calculated using the 2^-ΔΔCt^ method (Livak & Schmittgen, 2001) and were statistically compared using a two-sample t-test in Excel with XLSTAT (version 2020.3) (https://www.xlstat.com/en/).

### 2.5 Abundance of AmGSTF1

The analysis of B. sterilis GSTF1 levels was conducted using a total protein extract obtained from leaf tissue. The protein extraction and ELISA procedures were done according to (Torra et al., 2021). In brief, ∼85-90 mg of fresh leaf tissues were ground using liquid nitrogen, and then 900 μL of extraction buffer (100 mM Tris-HCl, 150 nM NaCl, 5 mM EDTA, 5% glycerol, 2% PVPP, and 10 mM DTT; pH 7.5) was added to the ground material. The mixture was subjected to ice incubation for approximately 10 mins and subsequently centrifuged at 12,000 ×g. The resulting supernatants were collected and subjected to a second centrifugation at the same speed for 15 mins. The total protein concentration was determined using the Bradford assay, following the manufacturer’s protocol (Bio-Rad Laboratories, United States). An ELISA was performed to measure the concentration of the polypeptides recognized by the anti-AmGSTF1-serum from *B. sterilis* protein samples, following the methods described by Torra et al., 2021.

### 2.6 Uptake and translocation studies

The adaxial surface of the second leaf from both the R and S plants was treated with [^14^C]pyroxsulam solutions mixed with the pyroxsulam commercial formulation. For this purpose, a micro-applicator PB-600 (Hamilton Company, Reno NV, USA) was used. The herbicide application rate was 18.75 g ai ha^−1^ in a 250 L ha^−1^ spraying volume for pyroxsulam, and the specific activity of the solution was 0.54 kBq μL^−1^. The plants were kept in a growth chamber under specified growing conditions, and the evaluation was done at 12 and 120 hours after treatment (HAT). [^14^C]pyroxsulam penetration and translocation studies were done as described by Palma-Bautista *et al*., 2023. The non-absorbed radiolabeled pyroxsulam’s radioactivity was analyzed using liquid scintillation spectrometry (LSS), and the absorption and translocation rates were calculated according to Silva *et al*., 2020. The percentage of ^14^C-herbicide recovery was greater than 85% in all samples assayed.

### 2.7 Antioxidant enzyme activity

Approximately 1g of leaf material was utilized for the assays of antioxidant enzyme activity. The plant materials were homogenized in 3 ml of ice-cold 100 mm K-phosphate buffer (pH 6.8) containing 0.1 mm EDTA for 5 min and filtered using a cheesecloth. Subsequently, the homogenate was centrifuged at 16,000 g for 15 min, and the resulting supernatant was used for the enzyme activity experiments. All the steps were done at a temperature of 0–4°C. The catalase, peroxidase, and superoxide dismutase (SOD) activities were done following the methodology described by Cavalcanti *et al*., 2004.

### 2.8 RNA-seq transcriptome and qRT-PCR validation of the selected contigs

Two control (S_C and R_C) and two treated (S_T and R_T) samples were used for the RNA-seq experiments. Total RNA was extracted from fresh leaf tissues, ±80-90Lmg per sample, using the RNeasy Mini Kit (QIAGEN) using the pre-optimized protocol (Sen *et al*., 2021). The integrity of the purified total RNA was assessed by agarose gel electrophoresis, and the concentration was measured using the Qubit 2.0 Fluorometer (Thermo Fisher Scientific) with the Qubit™ RNA HS Assay Kit (Invitrogen) before sending it for sequencing. The sequencing was performed at Novogene, China, using the Novaseq 6000 platform. Only samples with an integrity number greater than 7 and a concentration of ≥ 20 ng/uL were further selected for next-generation sequencing. Each treatment had five biological replicates, and the transcriptome sequencing parameters included 150 bp pair-end reads with a minimum of 30 million reads per sample. The clean raw reads obtained after quality control, including demultiplexing, trimming, and adaptor removal, were de-novo assembled using OmicsBox software (version 1.4.11). The assembled reads were then mapped back to the reference backbone generated by the de-novo assembly using the TRINITY program (Grabherr *et al*., 2011; Langmead & Salzberg, 2012). Unique mapped reads were considered for the gene expression quantification. The read counts were Log2 transformed and normalized using the scaling method. Further, biases in the sequence datasets and differences in transcript sizes were corrected using the RPKM algorithm to obtain accurate estimates of relative expression levels. The transcript levels of unigenes were calculated using the RSEM software package in OmicsBox (ver. 1.4.11) (Langmead & Salzberg, 2012; Li & Dewey, 2011). The assembly completeness was assessed using BUSCO (Seppey *et al*., 2019), and the coding region prediction was completed using Transdecoder (Haas *et al*., 2013). Initially, the transcript sequences were blasted against the non-redundant protein sequences in NCBI using the BLASTX-fast search program with an e-value cut-off of less than 1.0E^-5^. The Blast2GO platform under the functional analysis module within OmicsBox (ver 1.4.11) was used for this purpose. Gene ontology (GO) annotation was carried out in Blast2GO, utilizing the GO database (Consortium, 2004) and metabolic pathways from the Kyoto Encyclopedia of Genes and Genomics (KEGG) (Kanehisa *et al*., 2016), EggNOG, and orthologous groups (COG) (Huerta-Cepas *et al*., 2019). Candidate genes related to metabolism were selected based on those predicted to be up-regulated in both R_C vs S_C and R_T vs S_T comparisons (Wrzesińska-Krupa *et al*., 2023). These genes were further validated by qRT-PCR, as described in section 2.4. In this research, the P450s, GSTs, GTs, and ABC transporter genes that exhibited at least a 2-fold statistically significant change in at least one out of the four comparisons between treatments (R_C vs S_C, R_C vs R_T, S_T vs S_C, and R_T vs S_T) were validated (Wrzesińska-Krupa *et al*., 2023). The list of primers used for qRT-PCR is shown in supplementary table 1.

## 3. Results

### 3.1 Dose-response experiments and TSR mechanisms

#### 3.1.1 Sensitivity to pyroxsulam and cross-resistance to other herbicides

A series of herbicide experiments with pyroxsulam (with and without inhibitors) were conducted to determine the resistance level in the *B. sterilis* population from the CZ (Table 2 and Figures 1A and 1B). The pyroxsulam dose-response experiments revealed that the GR50 of the R and S biotypes are ∼371 (± 55.4) g a.i. ha^−1^ and ∼4 (± 3.8) g a.i. ha^−1^ (Table 2). Based on the RF, the R population was ∼86 times more resistant than the S population (Table 2). No significant growth reduction and/or visual damage were observed in *B. sterilis* treated with the recommended doses of malathion (1000Lg a.i. ha^−1^) and NBD-CI (270Lg a.i. ha^−1^). However, when malathion and NBD-CI were applied prior to pyroxsulam, there was a significant decrease in the RF compared to the plants treated with pyroxsulam alone (∼2X and ∼1.2X, respectively; Table 2). These results indicate the involvement of metabolic resistance in pyroxylam resistance. In addition to pyroxsulam, the R biotype from the CZ was also found to be cross-resistant to selected ALS, acetyl CoA caraboxylase (ACCase)-inhibiting, and pre-emergent herbicides, including propoxycarbazone-sodium, sulsulfuron, pinoxaden, propoxycarbazone plus mesosulfuron, pinoxaden plus pyroxsulam, flufenacet plus diflufenican, and halauxifen plus pyroxsulam plus florasulam (Figure 2).

**Figure 1:**
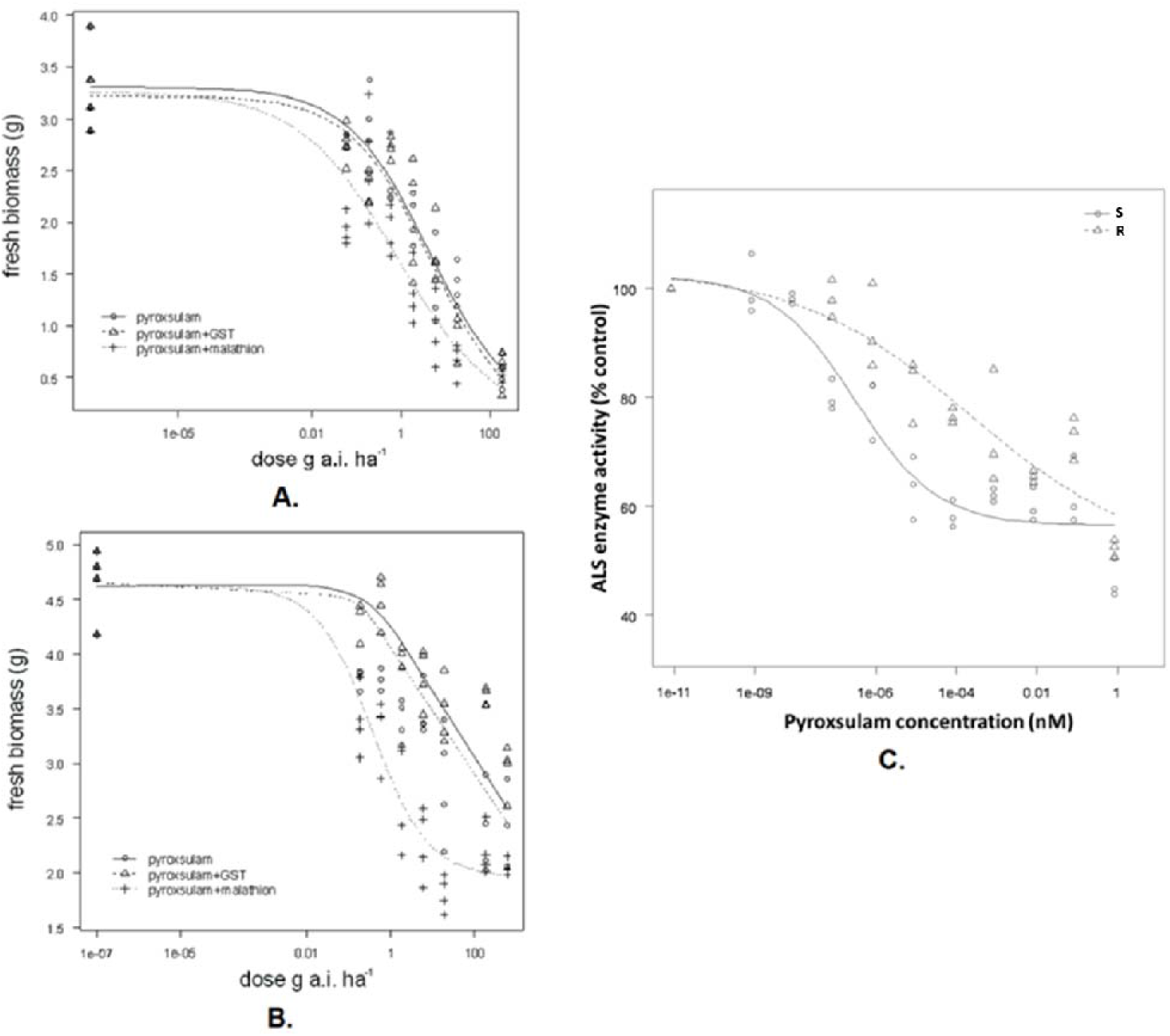
ALS dose response experiments and enzyme activity analysis. Results for fitted logarithmic dose–response curves with pyroxsulam for *B. sterilis* **(A)** susceptible (S) biotype, **(B)** the resistant (R) biotype. (**C.)** Results for fitted logarithmic dose–response curves with ALS enzyme activity for *B. sterilis*.

**Figure 2:**
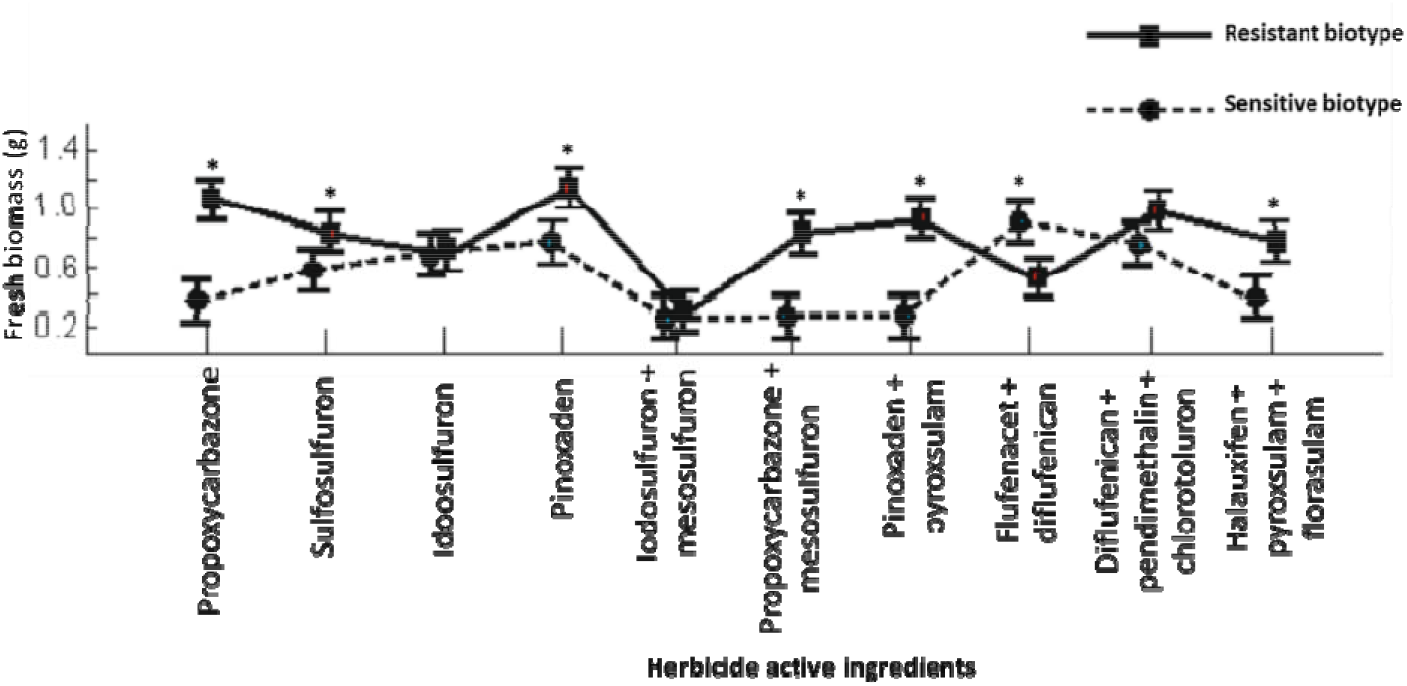
Results for cross-resistance experiments. The “*” indicates significant at the 5% significance level.

**Table 2:** Result from the non-linear regression analysis of pyroxsulam dose–response experiment for resistant (R) and susceptible (S) biotype. In the table ‘d’LisLthe upper limit, ‘b’LisLthe slope around the GR50, ‘GR50’LisLthe rate of herbicide (g a.i. ha^−1^) required to reduce above ground biomass weight by 50%, ‘SE’ isLthe standard error.

| Biotype | b (SE) | d (SE) | GR50 (SE) | Resistance factor |
| --- | --- | --- | --- | --- |
| R_ pyroxsulam | 19.88 (5.49) | 10.06 (5.53) | 370.55 (55.4) | 86.11 |
| S_ pyroxsulam | 11.04 (3.19) | 11.01 (6.21) | 4.30 (3.8) |  |
| R_ pyroxsulam + malathion | 18.66 (6.23) | 11.82 (2.89) | 36.60 (13.3) | 44.58 |
| S_ pyroxsulam + malathion | 24.19 (6.69) | 10.77 (4.22) | 0.82 (0.7) |  |
| R_ pyroxsulam + NBD-Cl | 14.63 (4.11) | 11.10 (6.77) | 356.48 (22.4) | 74.42 |
| S_ pyroxsulam + NBD-Cl | 21.81 (6.13) | 10.77 (4.22) | 4.79 (3.5) |  |

#### 3.1.2 ALS enzyme activity

To further confirm the resistance to pyroxulam in the R biotype, we determined the inhibition of ALS enzyme activity compared between the S and R biotypes. We rationalized that it would require a higher concentration of pyroxulam to inhibit ALS activity in the R biotype. In the absence of pyroxulam, similar ALS activity was observed in both the R and S biotypes. As expected, we found a significant difference in ALS activity between the R and S biotypes in the presence of pyroxulam (Figure 1C). The IC50 of the S biotype was 3.0978e^-07^, while the IC50 of the R biotype was 2.2583e^-04^. Using these IC50s, we calculated the RF (∼729) and confirmed that the R biotype required a higher concentration of pyroxulam to inhibit ALS activity. Taken together, the ALS enzyme activity experiments corresponded with the resistance phenotype of each biotype.

#### 3.1.3 *ALS* gene sequencing and ploidy level variation analysis

To explore the underlying mechanisms that confer pyroxulam resistance in the R biotype, we first investigated the point mutation of the ALS protein. Approximately 30 plants that survived pyroxulam treatments from the S and R biotypes were used for ALS mutation analysis. The known ALS mutation positions (Ala-122, Pro-197, Ala-205, Phe-206, Asp-376, Arg-377, Trp- 574, Ser-653, and Gly-654) were analysed. We found no ALS mutations in the R biotype (data not shown). Therefore, we conclude that pyroxsulam resistance in the R biotype is not due to ALS mutations. Additionally, the flow cytometry analysis showed that there was no difference in DNA content between the S and R biotypes (Figure 3A). This result suggests that the variation in ploidy level does not involve pyroxsulam resistance in the R biotype.

**Figure 3:**
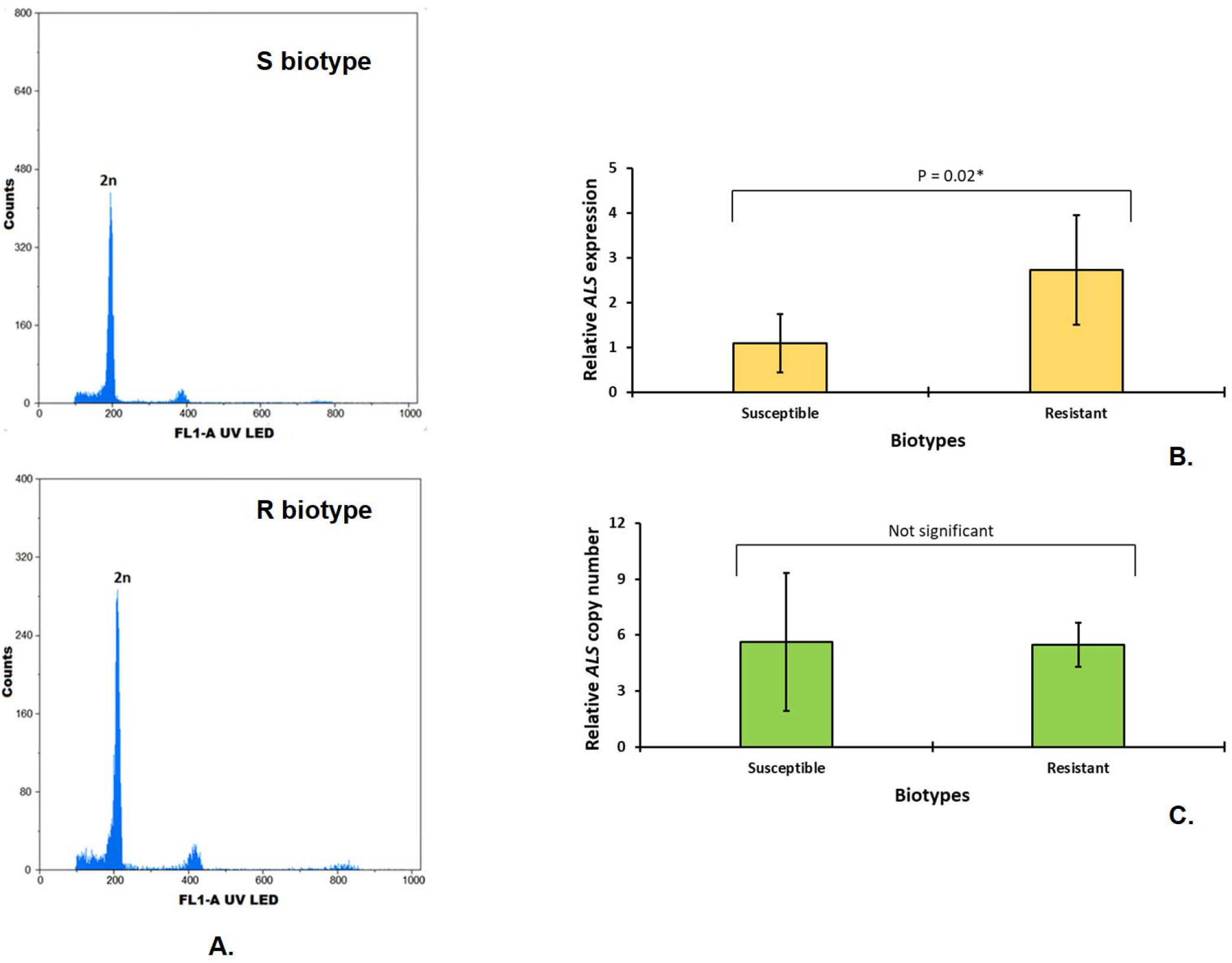
Flow cytometry and quantitative real-time PCR experiments. **(A.)** Flow cytometry profile of S and R biotype of *Bromus sterilis*, **(B.)** Relative *ALS* gene expression level and **(C.)** Relative *ALS* gene copy number. The results of quantitative real-time PCR experiments were compared by a two-sample t-test at 5% significance level. “*” and “NS” represents significant at 5% significance level and not significant, respectively.

#### 3.1.4 Relative *ALS* gene expression and copy number studies

An increased target gene copy number is another cause of target site resistance. Therefore, we explore the transcript expression of *ALS* in the S and R biotypes by qRT-PCR. It is interesting that the relative expression of the *ALS* in the R biotype was significantly (∼2.5-fold) higher than that of the S biotype (Figure 3B). For herbicide resistance, an increase in expression and gene copy number is a known mechanism to confer resistance to glyphosate in *Amanranthus* spp. (García *et al*., 2020). Since we observed an increased expression of *ALS* in the R biotype, we next determined the ALS copy number in the S and R biotypes. There was no significant difference in the relative *ALS* gene copy number between the R and S biotypes (Figure 3C). Collectively, our results suggest that a significant increase in expression of the *ALS* in R biotypes without an increase in *ALS* copy number might confer resistance to pyroxulam in *B. sterilis*.

### 3.2 Non-target site mechanisms

3.2.1 De novo assembly and functional annotation of the *B. sterilis* transcriptome

Our previous results showed that TSR is unlikely to be the main cause of significant enhanced resistance to pyroxsulam in the R biotype. Therefore, we next investigate the involvement of NTSR in mediating resistance through transcriptome analysis (RNA-seq). Due to the absence of transcriptome data for *B. sterilis* or its related species, de novo assembly was performed to create the reference transcriptome. The details of the de-novo assembly of the *B. sterilis* cDNA reference transcriptome are presented in Table 3. The overall results are represented using PCA plot and it shows samples clustered separately (supplementary figure S1). Total 183,770 transcripts were obtained with 77,259 total genes (N50-2,184 bp) (Table 3). The coding regions involved 65% complete sequences, 20% and 7% partial sequences of 5’ and 3’ respectively, and 8% internal sequences. BUSCO analysis showed 61.02% complete single copy, 1.84% complete duplicated, 10.05% fragmented, and 26.8% missing (Table 3). Among the total number of genes, 11,019 can be functionally annotated by Gene Ontology (GO) analysis. The functional annotation of the obtained sequences was done using EggNOG v6.0 (Hernández-Plaza et al., 2023). The majority of the genes were involved in “cellular processes and signaling” (6539 genes) and “metabolism” (5330 genes). A few others were implied for “information storage and processing” (4782 genes), while others remained poorly categorized (Figure 4). Among the cellular and signaling processes, our analysis predicted that the majority of the genes (∼83% from the 6539 genes) belonged to signal transduction mechanisms (∼44%), posttranslational modification, protein turnover, chaperones (∼29%) and intracellular trafficking, secretion, and vesicular transport (∼10%). Among the 5330 metabolism-related genes, most of the genes were involved in carbohydrate transport and metabolism (∼20%), amino acid transport and metabolism (∼18%), and secondary metabolite biosynthesis, transport, and catabolism (∼17%). Concerning the 4782 information storage and processing genes, the gene functions varied from replication, recombination, and repair to chromatin structure and dynamics (Figure 4).

**Figure 4:**
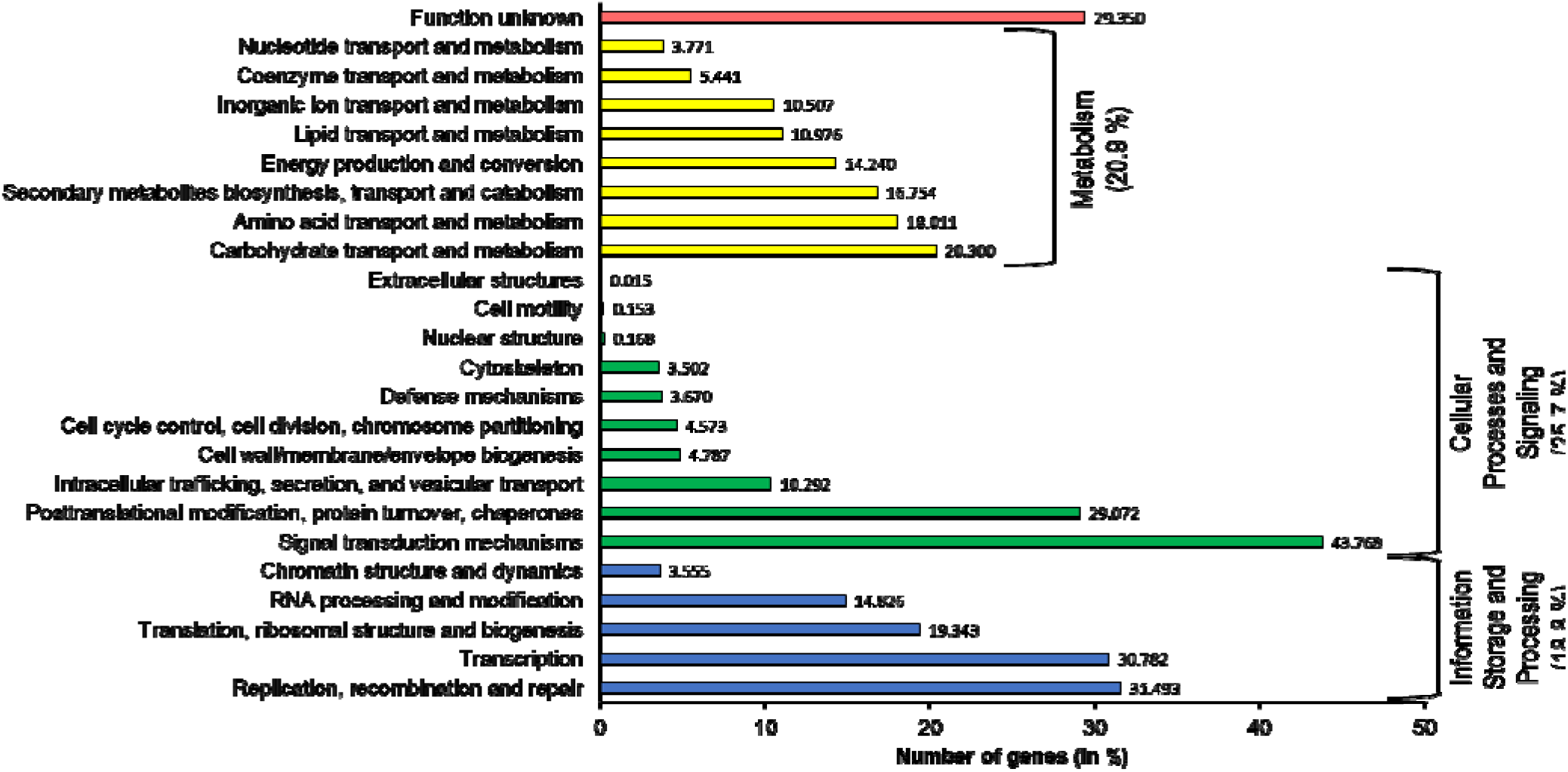
Functional annotation of the *B. sterilis* transcriptome. 11019 out of 77,259 gene were successfully annotated for gene ontology (GO) using EggNOG v6.0. Among these 11019 genes, 29.35% had some unknown function, while the others were categorized into three GO groups (information storage & processing, cellular processes & signalling and metabolism).

**Table 3:** Results from de-novo assembly of *Bromus sterilis* cDNA reference transcriptome.

|  | cDNA reference transcriptome |
| --- | --- |
| Total transcripts | 183,770 |
| Total genes | 77,259 |
| Percent GC | 47.87 |
| Total assembled bases (all transcripts) | 279,171,412 |
| Total assembled bases (longest isoform per gene) | 91,817,672 |
| N50 contig size | 2,184 bp |
| Median contig size | 1,128 bp |
| Average contig size | 1,519.13 bp |
| No. contigs with EggNOG annotation | 11,019 |
| Number of GO annotations | 78,928 |

| Coding Regions Summary [super transcripts] |  |
| --- | --- |
| **Complete** | 36,946 (65%) |
| **5' partial** | 11,233 (20%) |
| **3' partial** | 3,955 (7%) |
| **Internal** | 4,441 (8%) |
| BUSCO results |  |
| **Complete single copy** | 3,002 (61.02%) |
| **Complete duplicated** | 90 (1.84%) |
| **Fragmented** | 492 (10.05%) |
| **Missing** | 1,312 (26.8%) |

#### 3.2.2 Differential gene expression and functional analysis

To gain more information on NTSR mechanisms, we performed differentially expressed genes (DEGs) analysis between the untreated R and S biotypes. Additionally, we also performed DEG analysis between pyroxsulam-treated and untreated biotypes. The comparison between untreated and pyroxulam-treated samples will provide information on the differential effects of herbicide treatment in the R and S biotypes. The list of genes differentially expressed in R and S was presented in supplementary tables S2-S5. For the untreated plants, 2706 and 2501 genes were up- regulated and down-regulated, respectively, in the R biotype. Differential expression between herbicide-treated R and S plants (R_T vs S_T) identified 1786 genes that were up-regulated and 1663 genes that were down-regulated in the R plants. The comparison between S_T and S_C plants indicated 3017 up-regulated and 2274 down-regulated genes in the S_T plants relative to S_C. On the other hand, the comparison of transcript expression between herbicide-treated and untreated R plants identified 2979 up-regulated and 1300 down-regulated genes in the R_T plants relative to R_C. The full details on the number of differentially expressed genes for each experimental condition are shown in supplementary figure S2. The heatmaps and the volcano plots for the individual comparative analyses are shown in the supplementary figures S3–S6. To identify the functions of candidate genes that confer NTSR in the R biotype, gene set enrichment analysis (GSEA)-based bar plots for each analysis were done (supplementary figures S7–S10). In all the comparisons, GSEA predicted that the highest number of DEGS were associated with vesicle-mediated transport. In the cases of R_C vs S_C and R_T vs S_T, vesicle-mediated transport was followed by intercellular organelle lumen, nuclear lumen, organelle lumen, membrane-enclosed lumen, and endosomes. In contrast, for S_C vs S_T, the vesicle-mediated transport was followed by cytoskeletal protein binding, cytoskeleton, transmembrane transport, and nuclear transport. For R_C vs R_T, vesicle-mediated transport was followed by cytoskeletal protein binding, intercellular organelle lumen, membrane-enclosed lumen, nuclear lumen, and organelle lumen.

#### 3.2.3 Candidate herbicide-metabolic resistance contigs selection and validation

Enhanced herbicide detoxification is a well-studied NTSR mechanism in various dicot and monocot weed species [9]. Furthermore, the reduced resistance in R biotypes when pre-treated with inhibitors for herbicide detoxification enzyme families (Table 2) indicated an important role of herbicide detoxification on resistance phenotype. To identify candidate genes that function in herbicide detoxification, we compared the genes that were up-regulated in both the R_C vs S_C and R_T vs S_T data sets. Based on these criteria, we identified 57 transcripts as putative candidate genes that might confer resistance to pyroxsulam in *B. sterilis* (Table 4). We were able to annotate 19 contigs for CytP450s (5 transcripts), GSTs (5 transcripts), GTs (6 transcripts), and ABC transporters (3 transcripts). The remaining 38 transcripts were annotated to other families, such as other transporter proteins, transcription factors, reductases, disease-resistance proteins, peroxidases, and heat-shock proteins. In the scope of this study, we only validated the expression of CyP450s, GSTs, GTs, and ABC transporter genes (which had shown at least 2-fold statistically significant changes in at least one out of four comparisons between treatments) by qPCR. 12 contigs showed significantly higher transcript expression levels in the R samples than those of the S samples (Table 5). Collectively, a significant increase in transcript expression of genes encoded by multiple detoxification enzyme families corresponded with increased resistance to pyroxsulam as well as the cross-resistance to other herbicides observed in the R biotype.

**Table 4:** List of the prime candidates for metabolic-resistance genes in *Bromus sterilis* by RNA-Seq.

| Gene name |  | Gene family | R_T and<br>S_T<br>(Fold<br>change) | R_C and<br>S_C<br>(Fold<br>change) |
| --- | --- | --- | --- | --- |
| TRINITY_DN65097_c0_g1 | *Cytochrome P450 711A1* | Cytochrome P450s | 3.195 | 4.031 |
| TRINITY_DN1068_c3_g1 | *Cytochrome P450 71A1* |  | 2.027 | 1.140 |
| TRINITY_DN2299_c0_g1 | *Cytochrome P450 76M5* |  | 1.480 | 1.651 |
| TRINITY_DN5260_c0_g1 | *Cytochrome P450 72A15* |  | 1.337 | 1.805 |
| TRINITY_DN16753_c0_g1 | *Cytochrome P450 CYP709C1* |  | 1.190 | 3.890 |
| TRINITY_DN13721_c2_g1 | *Glutathione S-transferase lambda2*<br>*(GSTl2)* | Glutathione S-transferases | 1.410 | 1.506 |
| TRINITY_DN2559_c1_g3 | *Glutathione S-transferase U17* |  | 1.735 | 3.508 |
| TRINITY_DN1352_c1_g2 | *Glutathione S-transferase 4* |  | 1.514 | 4.345 |
| TRINITY_DN2115_c0_g3 | *Glutathione S-transferase GSTU6* |  | 1.306 | 1.910 |
| TRINITY_DN2838_c1_g2 | *Glutathione S-transferase T2* |  | 1.001 | 2.324 |
| TRINITY_DN1485_c0_g2 | *UDP-glycosyltransferase 73C5* | Glycosyltransferases | 2.136 | 2.406 |
| TRINITY_DN3596_c0 | *UDP-glycosyltransferase 75C1* |  | 1.541 | 4.575 |
| TRINITY_DN2821_c0_g4 | *UDP-glycosyltransferase 88F5* |  | 1.263 | 2.390 |
| TRINITY_DN4889_c0_g1 | *UDP-glycosyltransferase 87A2* |  | 1.146 | 1.661 |
| TRINITY_DN7848_c0_g6 | *UDP-glycosyltransferase 74E1* |  | 1.047 | 3.828 |
| TRINITY_DN408_c1_g2 | *UDP-glycosyltransferase 83A1* |  | 1.025 | 1.340 |
| TRINITY_DN20226_c0_g2 | *ABC transporter B family member 4* | ABC transporters | 2.330 | 8.706 |
| TRINITY_DN8286_c0_g3 | *ABC transporter G family member 41* |  | 1.178 | 1.218 |
| TRINITY_DN2655_c1_g1 | *ABC transporter G family member 11* |  | 1.076 | 1.074 |
| TRINITY_DN18985_c0_g1 | *Sulfate transporter 3.3* | Other transporters | 7.755 | 8.113 |
| TRINITY_DN40756_c0_g1 | *Bidirectional sugar transporter*<br>*SWEET13* |  | 6.222 | 6.254 |
| TRINITY_DN1158_c1_g1 | *Boron transporter 4* |  | 1.820 | 1.090 |
| TRINITY_DN3608_c0_g1 | *Zinc transporter 10* |  | 1.536 | 1.595 |
| TRINITY_DN6099_c0_g1 | *Nuclear transcription factor Y* | Transcription factors | 9.350 | 10.311 |
| TRINITY_DN31425_c0_g1 | *Transcription factor bHLH35* |  | 3.678 | 5.505 |
| TRINITY_DN5024_c0_g1 | *Transcription factor GLK1* |  | 3.078 | 2.974 |
| TRINITY_DN4669_c0_g1 | *Transcription factor NAI1* |  | 2.797 | 1.904 |
| TRINITY_DN2526_c2_g1 | *Transcription factor phytochrome*<br>*interacting factor-like 13* |  | 2.396 | 1.304 |
| TRINITY_DN10307_c1_g1 | *MYB13 transcription factor (MYB13-*<br>*1)* |  | 2.378 | 2.490 |
| TRINITY_DN3287_c0_g2 | *Lks2 gene for putative short*<br>*internodes family transcription factor* |  | 1.922 | 3.798 |
| TRINITY_DN8596_c0_g1 | *Transcription factor bHLH87* |  | 1.922 | 1.332 |
| TRINITY_DN9286_c1_g2 | *Transcription factor HHO5* |  | 1.391 | 2.245 |
| TRINITY_DN10047_c0_g1 | *Heat stress transcription factor A-2b* |  | 1.053 | 2.147 |
| TRINITY_DN949_c3_g4 | *NAC transcription factor (nac020*<br>*gene)* |  | 1.018 | 3.437 |
| TRINITY_DN17474_c0_g1 | *Ferredoxin--NADP reductase* | Reductases | 5.027 | 10.808 |
| TRINITY_DN7079_c1_g3 | *12-oxophytodienoate reductase 5* |  | 3.717 | 1.943 |
| TRINITY_DN10519_c1_g1 | *Aldo-keto reductase 3* |  | 3.380 | 3.013 |
| TRINITY_DN25284_c0_g1 | *2-alkenal reductase (NADP(+)-dependent)-like* |  | 3.185 | 8.704 |
| TRINITY_DN44814_c0_g1 | *Tropinone reductase homolog At5g06060-like* |  | 2.000 | 4.861 |
| TRINITY_DN5335_c0_g1 | *12-oxophytodienoate reductase 1-like* |  | 1.735 | 3.436 |
| TRINITY_DN1465_c0_g2 | *12-oxophytodienoate reductase 11* |  | 1.515 | 2.547 |
| TRINITY_DN1731_c0_g1 | *NAD(P)H-quinone oxidoreductase subunit O* |  | 1.109 | 1.101 |
| TRINITY_DN420_c2_g4 | *Disease resistance RPP13-like protein 3* | Disease resistance proteins | 10.708 | 10.361 |
| TRINITY_DN4956_c1_g1 | *Disease resistance protein RPM1-like* |  | 9.642 | 9.706 |
| TRINITY_DN378_c1_g2 | *Disease resistance protein Pik-2-like* |  | 8.391 | 8.953 |
| TRINITY_DN5409_c0_g1 | *Disease resistance protein RGA3* |  | 6.420 | 7.467 |
| TRINITY_DN8126_c0_g1 | *Disease resistance protein (Lr1)* |  | 6.343 | 8.344 |
| TRINITY_DN6574_c0_g1 | *Disease resistance protein RGA5-like* |  | 5.536 | 6.848 |
| TRINITY_DN5491_c0_g1 | *Disease resistance protein At3g14460* |  | 4.115 | 5.301 |
| TRINITY_DN4424_c0_g1 | *Disease resistance protein RGA2-like* |  | 3.093 | 1.433 |
| TRINITY_DN6669_c0_g1 | *Disease resistance protein RGA5-like* |  | 2.974 | 4.844 |
| TRINITY_DN3482_c1_g1 | *Disease resistance RPP13-like protein 3* |  | 2.432 | 1.520 |
| TRINITY_DN12404_c0_g1 | *Disease resistance protein RGA1* |  | 1.845 | 1.345 |
| TRINITY_DN844_c0_g1 | *Disease resistance protein At1g61300* |  | 1.306 | 1.788 |
| TRINITY_DN223_c0_g1 | *Disease resistance protein RGA1* |  | 1.290 | 1.003 |
| TRINITY_DN21041_c0_g1 | *Phospholipid hydroperoxide glutathione peroxidase* | Peroxidase | 10.077 | 11.402 |
| TRINITY_DN61846_c0_g1 | *Small heat shock protein* | Heat shock proteins | 2.035 | 1.904 |

**Table 5:** Validation studies with qRT-PCR. In this table, we validated the *P450s*, *GSTs*, *GTs* and *ABC transporter* genes which had shown statistically significant changes in R_T vs S_T and the qRT_PCR (fold change) denotes the ratio of gene expression values (calculated via 2^–ΔΔCt^ method) of R and S. In both the cases “*” represents significant at 5% significance level.

| Gene name | Gene name | RNA-seq (fold change) | qRT_PCR (fold change) |
| --- | --- | --- | --- |
| TRINITY_DN65097_c0_g1 | *Cytochrome P450 711A1* | 9.155* | 12.345* |
| TRINITY_DN1068_c3_g1 | *Cytochrome P450 71A1* | 4.074* | 8.073* |
| TRINITY_DN16753_c0_g1 | *Cytochrome P450*<br>*CYP709C1* | 2.281* | 6.865* |
| TRINITY_DN2559_c1_g3 | *Glutathione S-transferase*<br>*U17* | 3.330* | 5.869* |
| TRINITY_DN1352_c1_g2 | *Glutathione S-transferase*<br>*4* | 2.856* | 8.907* |
| TRINITY_DN2115_c0_g3 | *Glutathione S-transferase*<br>*GSTU6* | 2.473* | 9.876* |
| TRINITY_DN2838_c1_g2 | *Glutathione S-transferase*<br>*T2* | 2.001* | 8.008* |
| TRINITY_DN1485_c0_g2 | *UDP-glycosyltransferase*<br>*73C5* | 4.397* | 12.883* |
| TRINITY_DN3596_c0 | *UDP-glycosyltransferase*<br>*75C1* | 1.718* | 2.782* |
| TRINITY_DN2821_c0_g4 | *UDP-glycosyltransferase*<br>*88F5* | 2.400* | 6.557* |
| TRINITY_DN7848_c0_g6 | *UDP-glycosyltransferase*<br>*74E1* | 2.066* | 4.757* |
| TRINITY_DN20226_c0_g2 | *ABC transporter B family*<br>*member 4* | 5.029* | 9.871* |

#### 3.2.3 GSTF1: Enzyme-Linked Immunosorbent Assay (ELISA)

The protein extracts of S and R biotypes were analyzed by ELISA using anti-blackgrass GSTF1 (AmGSTF1) monoclonal antibody to confirm the immunoreactivity of the protein abundant and to determine its accumulation level. It has been reported previously that the orthologue protein of AmGSTF1 in *B. sterilis* was significantly increased in the herbicide-resistant *B. sterilis* population in the United Kingdom (Davis *et al*., 2020). AmGSTF1 is a well-defined biomarker for NTSR in blackgrass (Comont *et al*., 2020). Furthermore, increased transcript and protein expression of the orthologue AmGSTF1 have been reported in ryegrass (*Lolium rigidum*) and wild oat (*Avena fatua*) that showed NTSR phenotypes. Therefore, we tested whether the R biotype accumulated a higher level of AmGSTF1 ortholog protein than that in the S biotype. Western blotting analysis of crude protein extracts from the R and S biotypes showed that AmGSTF1-specific antibodies can detect the orthologue in B. sterlis. Using an ELISA assay, we demonstrated that the levels of AmGSTF1 immunoreactive polypeptides in the R biotype were significantly higher (∼5 times with P = 0.004) than those in the S population (Figure 5). According to Blastn analysis, a significant similarity of 71% identity (E-value = 1e^-48^) was observed between AmGSTF1 and TRINITY_DN1352_c1_g2.

**Figure 5:**
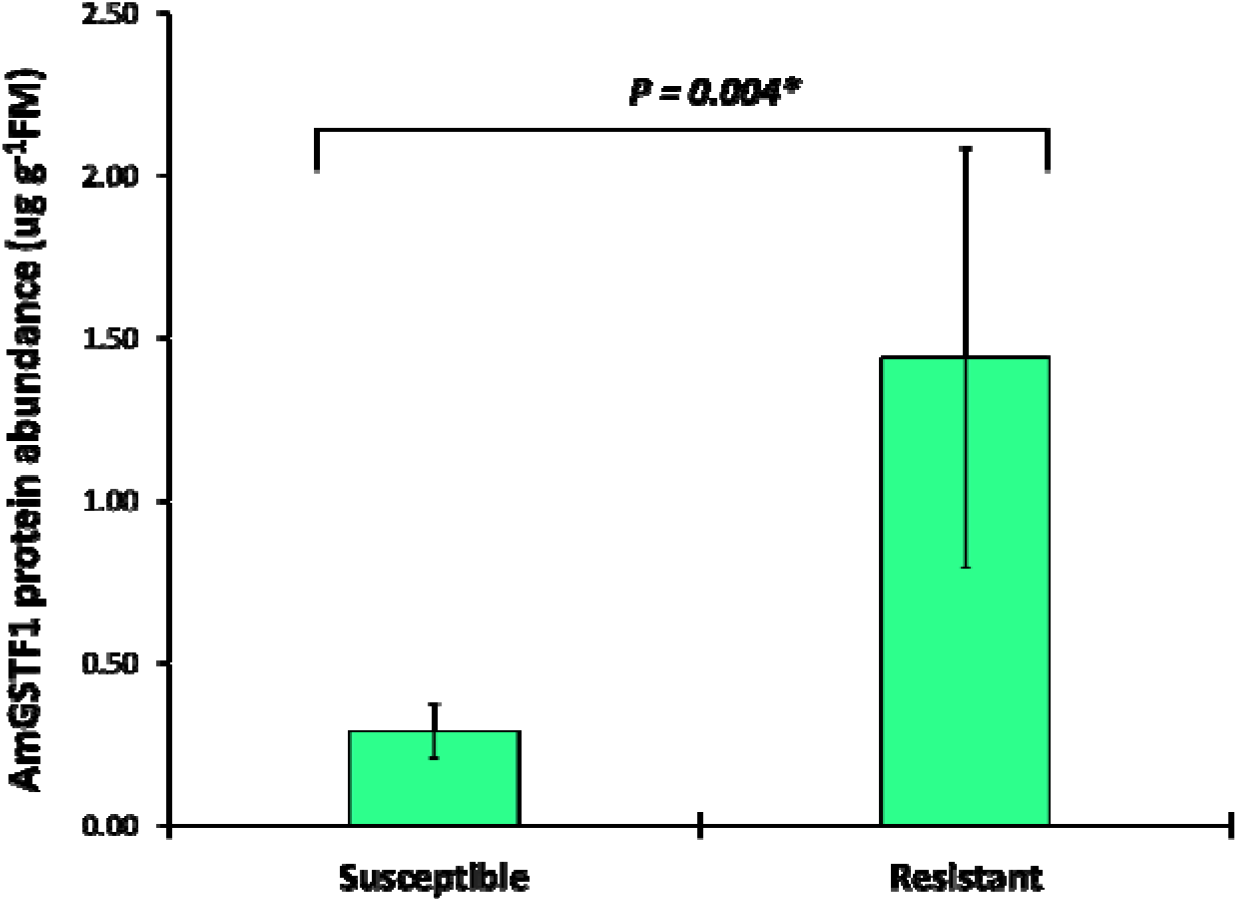
Analysis of the content of ortholog protein of AmGSTF1 (a biomarker protein) using Enzyme-Linked Immunosorbent Assay (ELISA) in a susceptible and resistant biotype of *Bromus sterilis*. The bars represent average values from five biological replicates and the ELISA results were compared by two sample t-test at 5% significance level. “*” represent significant at 5% significance level.

#### 3.2.4 Catalase, peroxidase and SOD activities

Apart from detoxification genes, we identified various genes that function in redox homeostasis that were up-regulated in the R biotype (Table 6). Since enzymatic antioxidant reactions are one of the main mechanisms of redox hemeostasis, we tested catalase, peroxidase, and superoxide dismutase (SOD) activities in R and S biotypes. The antioxidant enzyme activities were found highly significant (pL>L0.05) in the R biotype compared to the S biotype (Table 6). However, catalase (4 fold higher in R than in S) activity was highly abundant as compared to the other two antioxidant enzymes. The peroxidase and the SOD activity in R were found to be 1.4X and 2.1X higher than in S, respectively (Table 6). The overall increased enzymatic antioxidant reactions in the R biotype are positively correlated with increased transcript expression of redox-related genes.

**Table 6:** Results of antioxidant enzyme activity. The results were compared by two sample t- test at 5% significance level. The “*” shows a significant difference between resistant biotype (R) and the susceptible biotype (S).

| **Biotype** | **Catalase activity**<br>**(nmol min<sup>-1</sup> mg<sup>-1</sup>**<br>**protein)** | **Peroxidase activity**<br>**(nmol min<sup>-1</sup> mg<sup>-1</sup>**<br>**protein)** | **SOD activity**<br>**(U mg<sup>-1</sup> protein)** |
| --- | --- | --- | --- |
| S | 0.28 (±0.05) | 0.026 (± 0.006) | 0.017 (± 0.001) |
| R | 1.04 (± 0.35)* | 0.036 (± 0.001)* | 0.036 (± 0.002)* |

#### 3.2.5 [^14^C]pyroxsulam penetration and translocation in *B. sterilis*

The reduced absorption and translocation of herbicides are another NTSR mechanism that confers strong resistance to herbicides, including glyphosate (Pan *et al*., 2021). We identified various transporters up-regulated in the R biotype, which could suggest the prospective involvement of absorption and translocation in mediating resistance to pyroxsulam. Using radio- labeled pyroxsulam, we determined the absorption and translocation of pyroxsulam in the S and R biotypes (Table 7). The pyroxsulam absorption in the S biotype (∼1.2% to ∼2.6%) was comparable to that in the R biotype (∼1.1% to ∼2.7%). For translocation, there were no significant differences between the treated leaves and roots at 12 HAT. However, the R biotype showed a significantly lower translocation of pyroxsulam to systematic leaves when compared to the translocation in the S biotype. It is interesting that while there were no significant differences in translocation in treated and systematic leaves compared between S and R biotypes, the R biotype showed reduced translocation from treated leaves to systematic leaves and root tissues at 120 HAT.

**Table 7:**
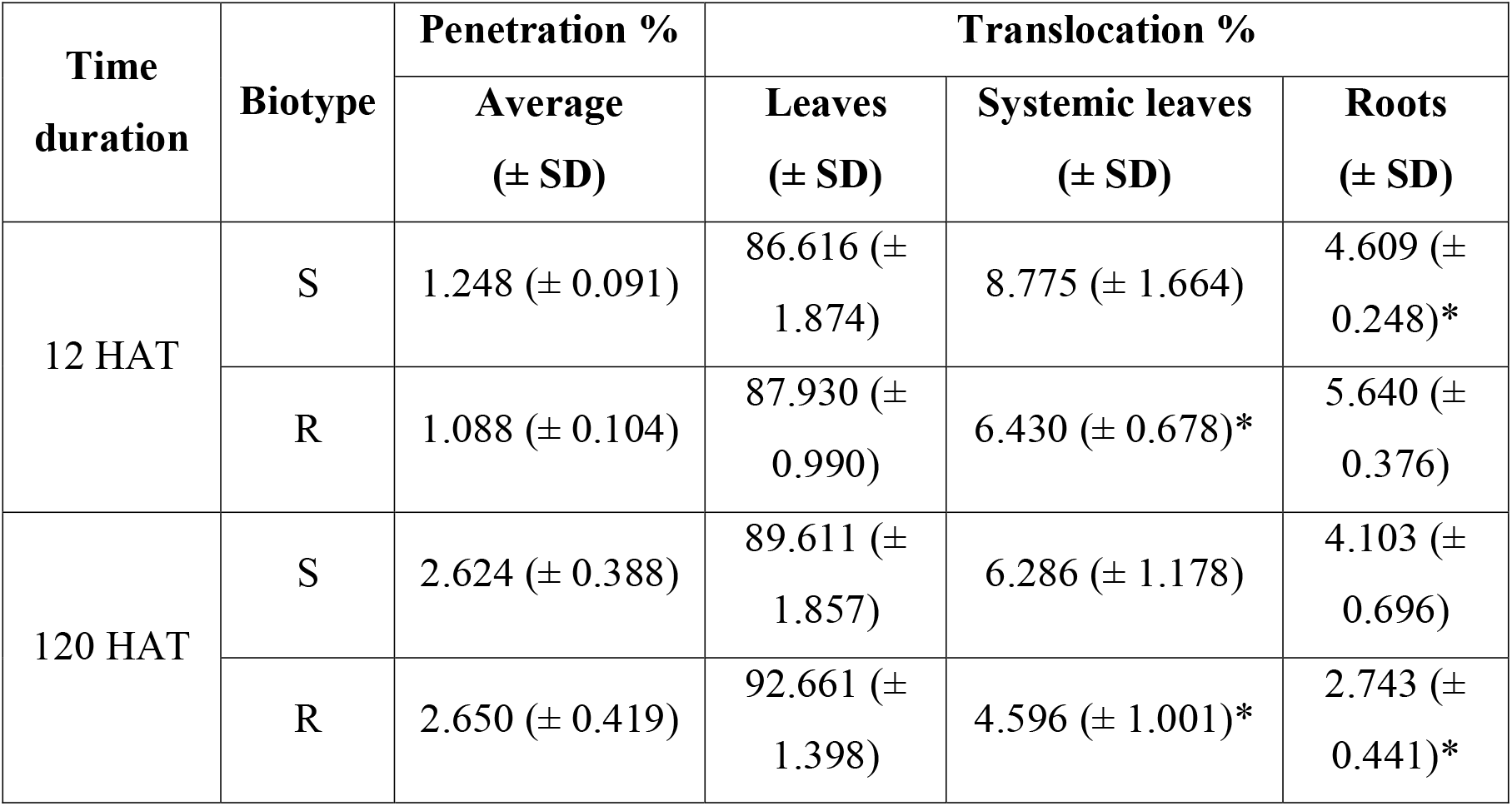
Herbicide penetration and translocation studies. The results were compared by a two-sample t-test at 5% significance level. The “*” shows a significant difference between the resistant biotype (R) and the susceptible biotype (S).

## 4. Discussion

The development of herbicide resistance in noxious weeds like *B. sterilis* is an obvious consequence of over-dependence on herbicides and is certainly a serious environmental issue. If left uncontrolled, herbicide-resistant weeds might also outcompete native plant species, leading to a loss of biodiversity in agricultural fields and stumbling effects on the entire ecosystem (Schütte *et al*., 2017). A detailed understanding of the mechanisms involved in resistance may provide novel approaches and new herbicides for weed control. The resistance mechanisms in herbicide-resistant weeds must be discovered before they become a significant challenge to sustainable agriculture, biodiversity conservation, and ecosystem health.

The objective of the current study was to unravel the molecular mechanisms underlying resistance to pyroxsulam in a *B. sterilis* biotype from the CZ. We present the novel reference transcriptome for this species in the study, which can serve as a valuable resource for understanding herbicide resistance in related *Bromus* species. The field-derived R biotype used in this study showed high levels of pyroxsulam resistance (RF ≈ 86), and interestingly, this resistance was reversed by inhibitors targeting herbicide detoxification enzyme families, namely Cyp450 and GSTs. Importantly, we confirmed the cross-resistance patterns to other post- and pre-emergent herbicides in this R biotype. We hypothesize that the cross-resistance to multiple herbicides might be due to the development of enhanced herbicide detoxification mechanisms (Yu & Powles, 2014; Reade *et al*., 2004). The multiple mutations in herbicide target proteins and the variation in ploidy level can confer cross-resistance to multiple herbicides. However, the lack of mutations and no polymorphisms in the ploidy level between the S and R biotypes indicate that these mechanisms do not involve in conferring resistant phenotype in the R biotype. It is intriguing that we observed a significant increase in the expression of the *ALS* gene in R biotype as compared to that in the S biotype. An increased expression of the herbicide target gene could confer resistance to a specific herbicide. There are several mechanisms that can contribute to gene overexpression, including gene amplification (Patterson *et al*., 2018), activating mutations within the regulatory regions (Prelich, 2012), or epigenetic activation (Chang *et al*., 2020). In this study, we focused solely on the *ALS* gene copy number, while other potential mechanisms remain to be explored. Our results showed a comparable copy number of ALS between the R and S biotypes, which suggests this mechanism is less likely to contribute to increased ALS expression in the R biotype. It is noteworthy that the association between gene copy number and herbicide resistance is a comparatively novel finding in herbicide resistance research and supported by a limited number of evidences (Patterson *et al*., 2018; Gaines *et al*., 2010). Therefore, additional experiments in a broader set of population needs to be done before drawing a final conclusion on the role of ALS copy number and herbicide resistance in *Bromus* spp.

Based on accumulated information, non-target site resistance (NTSR) has diverse molecular mechanisms, and enhanced herbicide detoxification and reduced herbicide absorption- translocation rates are the well-studied NTSR mechanisms. Herbicide detoxification typically involves three major phases. Phase I encompasses the conversion of the toxic herbicidal molecule into a more hydrophilic metabolite, enhancing its solubility for subsequent reactions. Thereafter, in phase II, the hydrophilic metabolite is conjugated with biomolecules such as glutathione or sugar. Finally, in phase III, the resulting metabolite undergoes transport to vacuoles or cells following additional reactions (Nandula *et al*., 2019). The major enzymes involved in Phase I are Cyp450 and their isoforms, while the major phase II and III enzymes include GSTs/GTs and ABC transporters, respectively (Gaines *et al*., 2020; Nandula *et al*., 2019). Based on RNA-seq analysis of R and S biotypes, we identified 57 main contigs as putative candidate genes, out of which 12 contigs (belonging to the CytP450, GST, GT and ABC transporter families) were validated via qRT-PCR. Five isoforms of CytP450 (TRINITY_DN65097_c0_g1, TRINITY_DN1068_c3_g1, TRINITY_DN2299_c0_g1, TRINITY_DN5260_c0_g1 and TRINITY_DN16753_c0_g1) were identified with significantly higher constitutive expression levels in the R plants than the S plants. The positive correlation between collective increased transcript expression of detoxification gene families and cross-resistance to multiple herbicides in the R biotype suggests that enhanced herbicide detoxification is one of the important mechanisms that protect R plants from herbicide damage.

Apart from herbicide detoxification, we also examined whether herbicide aborption- translocation could explain the resistant phenotype in the R biotype. The significant increased transcript expression of several transporters and a significant reduction in the translocation of pyroxsulam from treated leaves to systematic leaves in the R biotype suggest the prospective involvement of herbicide translocation in mediating herbicide resistance. There has been no reports of reduced translocation of pyroxsulam that confers resistance in any weed species. However, it has been shown that wheat has a lower translocate rate of pyroxulam than blackgrass, which likely links to the faster detoxification of this herbicide in wheat (deBoer et al., 2011). These mechanisms are the basis for pyroxsulam selectivity in wheat; hence, it might be possible that herbicide resistant grass weeds develop similar mechanisms to protect themselves against pyroxsulam toxicity. Therefore, it is important to further investigate this mechanism in other pyroxsulam-resistant weed species and monitor other brome populations for this herbicide resistance mechanism in Europe.

While detoxification and absorption/translocation are important to prevent the active pyroxsulam molecule from reaching its target site and causing damage, exposure to pyroxsulam could induce oxidative stress within the cell (Veith & Moorthy, 2018). In response to oxidative stress, the cell activates its antioxidant enzymes, such as catalase, peroxidases and SOD activities, to protect itself (Pérez-Torres et al., 2011). Intriguingly, the GO analysis revealed that various genes that function in redox homeostasis were significantly up-regulated in the R biotype. Indeed, we observed a significant increase in the activities of antioxidant enzymes of the R biotype compared to those of the S biotype. These results support the prospective involvement of redox homeostasis in herbicide resistance. Additionally, a previous study conducted on *Conyza bonariensis* found that glyphosate-induced stresses can enhance the activities of antioxidant enzymes in resistant plants (Piasecki *et al*., 2019). Therefore, future experiments to establish the link between redox homeostasis and herbicide resistance would provide an in-depth understanding of the role of signalling molecules such as reactive oxygen species in mediating NTSR. While AmGSTF1 has been identified as a functional biomarker of NTSR mechanisms through enhanced metabolism in *A. myosuroides* (Franco-Ortega *et al*., 2021) and *L. rigidum* (Torra *et al*., 2021), the biochemical functions of this protein remain elusive. AmGSTF1 has been proposed to function in the main regulatory network of NTSR since this protein has low glutathione conjugation activity (Cummins *et al*., 2013). Therefore, the detection of AmGSTF1 and its orthologue is considered a biomarker for NTSR but is unable to pinpoint the specific mechanisms (Onkokesung *et al*., 2022). In our study, we observed significant elevation of an orthologue of the AmGSTF1 proteins in the R biotype of *B. sterilis*, which corresponded with a cross-resistance pattern mediated by NTSR. These results suggest the important function of regulators such as AmGSTF1 in determining the outcome of herbicide resistance in plants. In fact, we identified the up-regulation of a number of transcription factors (TFs) in R biotypes. Future experiments, including knockout or overexpression of specific TFs, are needed to establish the role of TFs in mediating herbicide resistance.

In conclusion, our study reveals that *B. sterilis* populations from Central Europe (CZ) have developed high resistance to pyroxsulam, along with cross-resistance to other herbicide modes of action, via three distinct mechanisms, namely *ALS* gene overexpression, enhanced metabolism, and decreased translocation. The results from this study not only address the existing knowledge gap in the field but also lay the foundation for future research. Further investigations may involve integrated omics approaches, such as genomics-transcriptomics, transcriptomics-proteomics, or transcriptomics-proteomics-metabolomics, to gain a comprehensive understanding of the underlying molecular mechanisms and their interplay.

## Research funding

This work was supported by grant from the Long-term conceptual development of research organization [The Ministry of Education, Youth and Sports (Czech Republic)].

## Acknowledgements

The authors thank Novogene Co., Ltd. (Beijing, China) for sequencing and consultation on experimental design. JT gratefully acknowledges support from the Ministerio de Ciencia, Innovación y Universidades (Ramón y Cajal grant RYC2018-023866-I), Spain. AR is supported by “Excellent Team Grants” (2023-2024) from the Faculty of Forestry and Wood Sciences, Czech University of Life Sciences, Prague, Czechia. RB was supported by the Internal Grant Agency, (grant number 20233105) from the Faculty of Tropical AgriSciences, Czech University of Life Sciences in Prague.

## Competing interests

The authors declare that there are no conflicts of interest.

## Authorship contribution

**MKS**: overall project conceptualization, conducting experiments, overall result interpretation and data analysis, writing – original draft, writing – review & editing. **KH**: overall project conceptualization, data analysis, writing – review & editing. **NO**: conducting ELISA experiments, writing – review & editing. **JM**: conceptualization (penetration and translocation studies), data analysis, writing – review & editing. **JT**: conceptualization (penetration and translocation studies), data analysis, writing – review & editing. **PK**: methodology (greenhouse experiments), writing – review & editing. **GS**: methodology (protein extraction), writing – review & editing. **AG**: formal analysis (antioxidant enzyme assays), methodology, writing – review & editing. **RB**: formal analysis (flow cytometry), methodology, writing – review & editing. **VPS**: formal analysis (statistics), writing – review & editing. **AR**: RNA-seq data analysis, supervision, writing – review & editing. **JS**: overall project supervision, conceptualization, formal analysis, funding acquisition, writing – review & editing. All authors reviewed and edited the final manuscript.

## Availability of the sequence data

This Transcriptome Shotgun Assembly project has been deposited at NCBI under the BioProject: PRJNA991136. The version described in this paper is the first version. The BioProject and associated SRA metadata are available for the reveiwers at https://dataview.ncbi.nlm.nih.gov/object/PRJNA991136?reviewer=m2n6g996jrlacgdoapuhikivua in read-only format.

## Notes

### Competing Interest Statement

The authors have declared no competing interest.

