## Supplemental figures for "Transcriptomic response in pyroxsulam-resistant and susceptible *Bromus sterilis* identified three distinct mechanisms of resistance"

### Slide 1
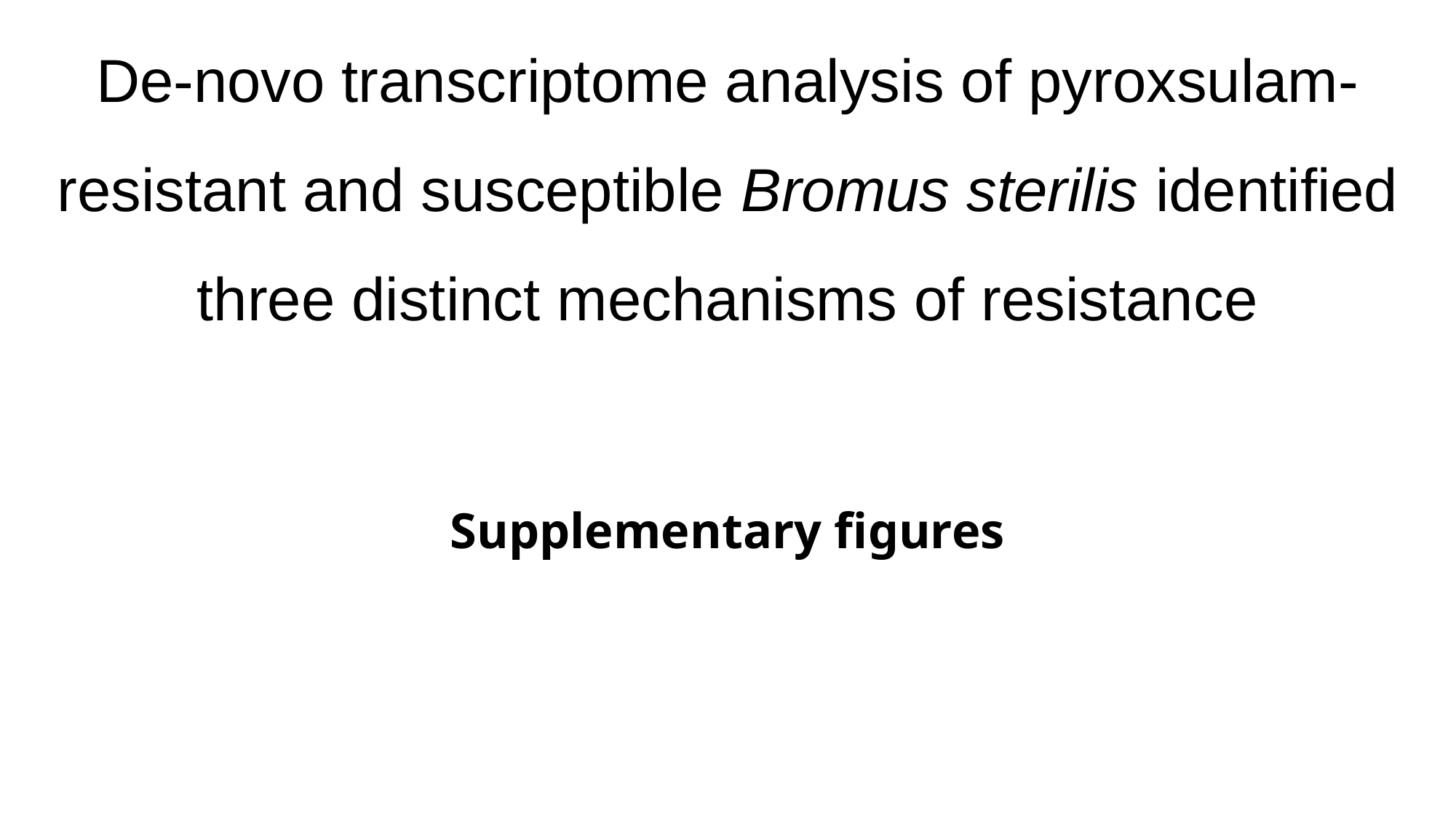

De-novo transcriptome analysis of pyroxsulam-resistant and susceptible Bromus sterilis identified three distinct mechanisms of resistance
Supplementary figures

### Slide 2
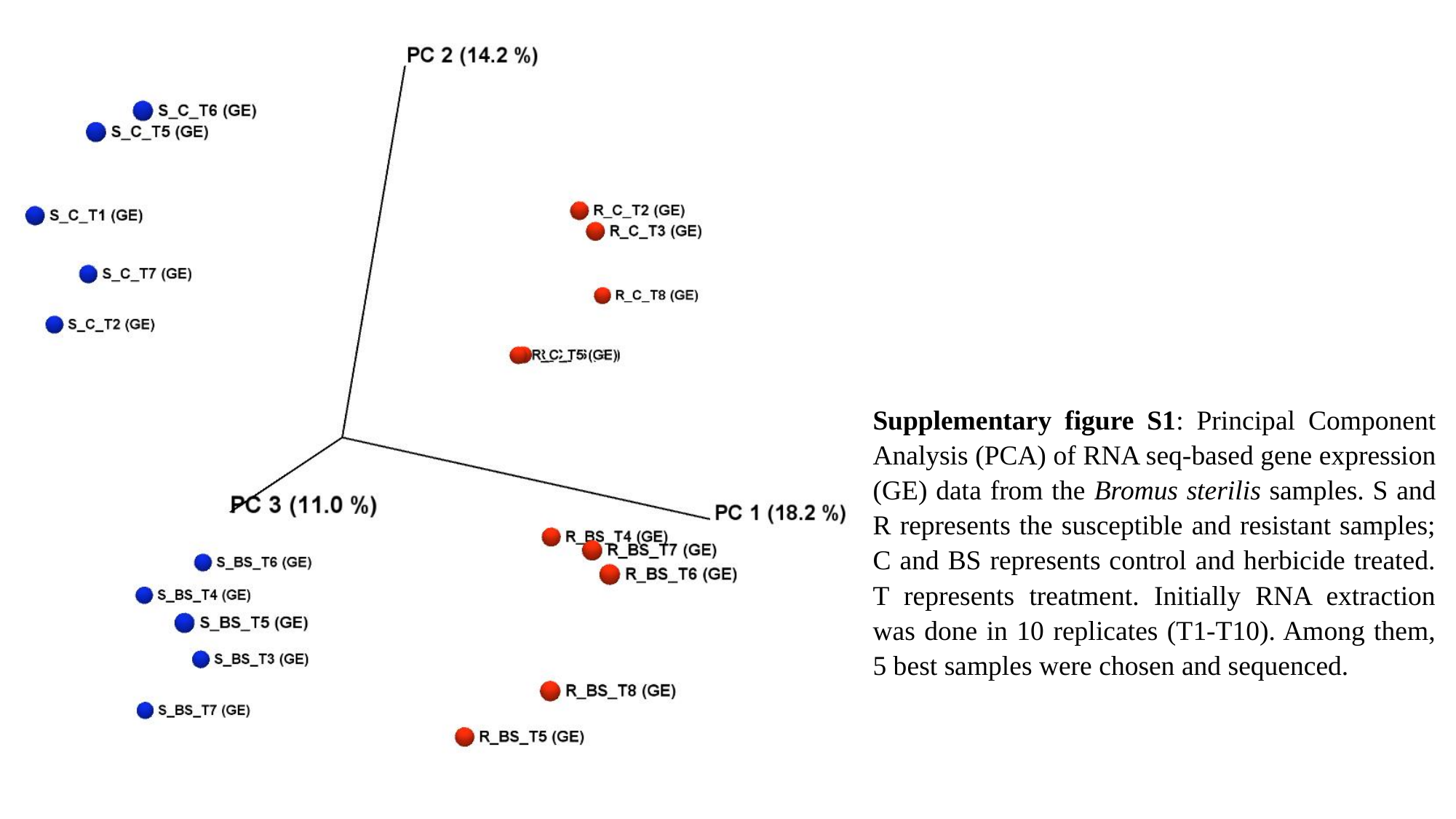

Supplementary figure S1: Principal Component Analysis (PCA) of RNA seq-based gene expression (GE) data from the Bromus sterilis samples. S and R represents the susceptible and resistant samples; C and BS represents control and herbicide treated. T represents treatment. Initially RNA extraction was done in 10 replicates (T1-T10). Among them, 5 best samples were chosen and sequenced.

### Slide 3
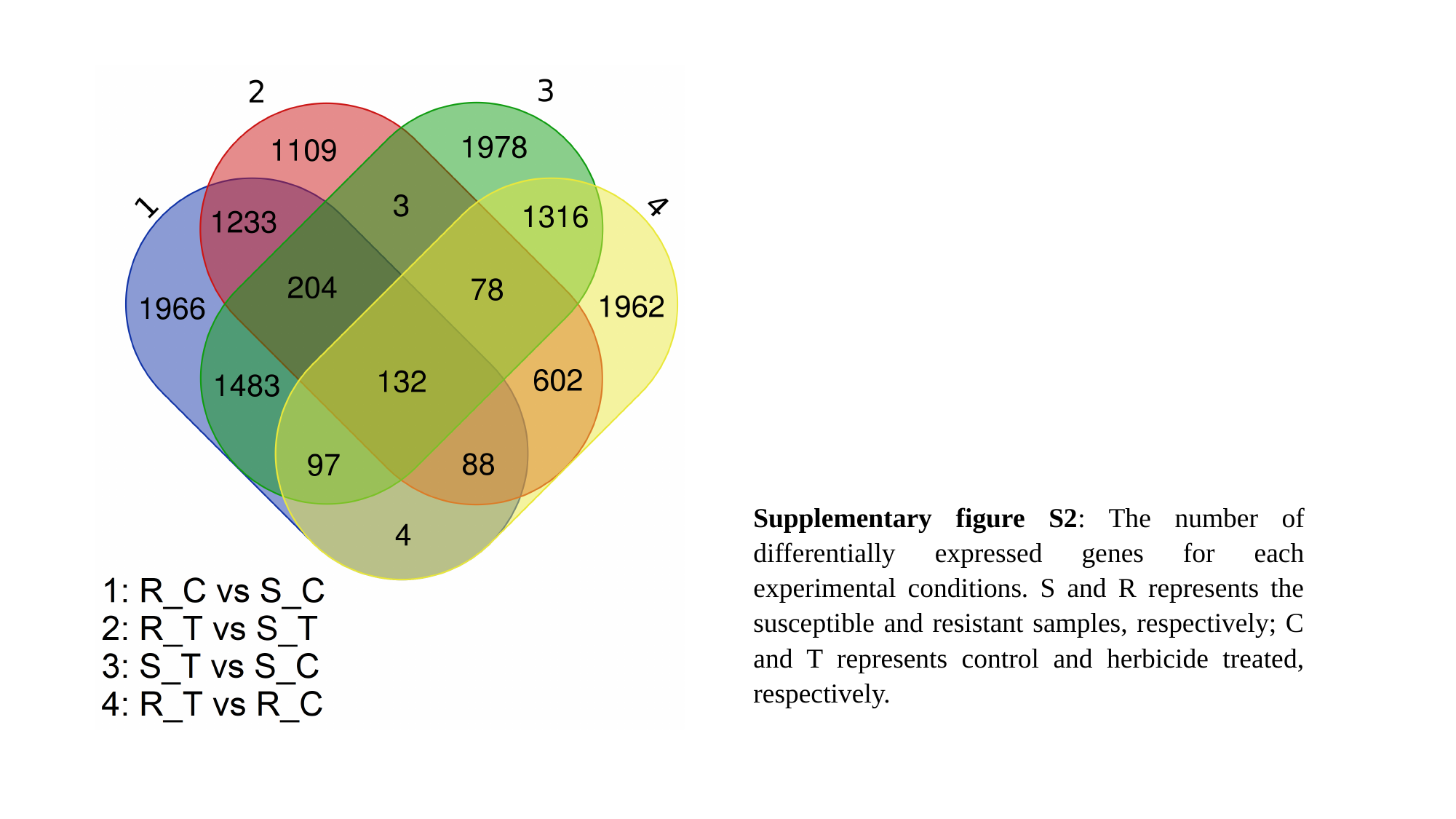

Supplementary figure S2: The number of differentially expressed genes for each experimental conditions. S and R represents the susceptible and resistant samples, respectively; C and T represents control and herbicide treated, respectively.

### Slide 4
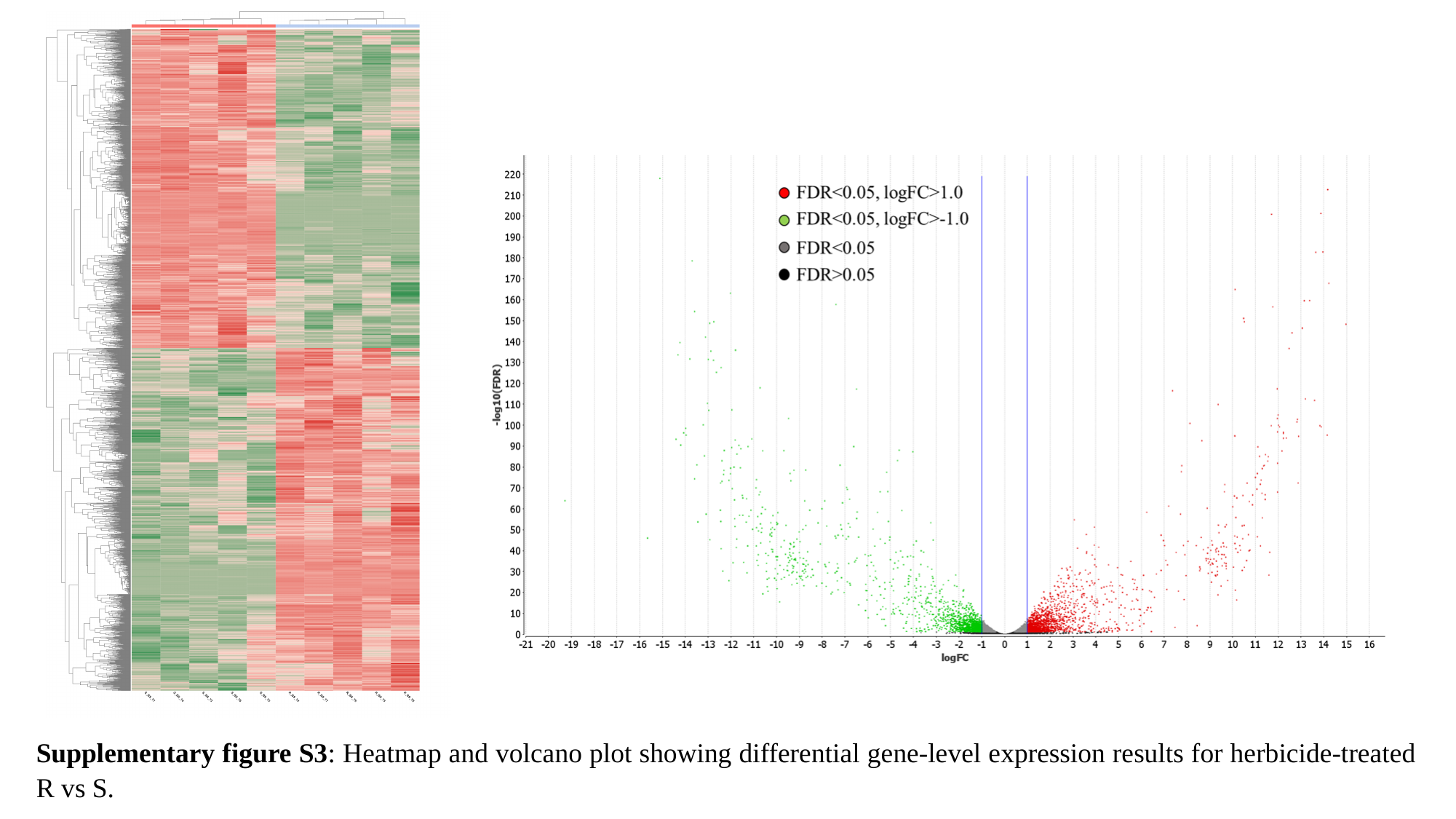

Supplementary figure S3: Heatmap and volcano plot showing differential gene-level expression results for herbicide-treated R vs S.

### Slide 5
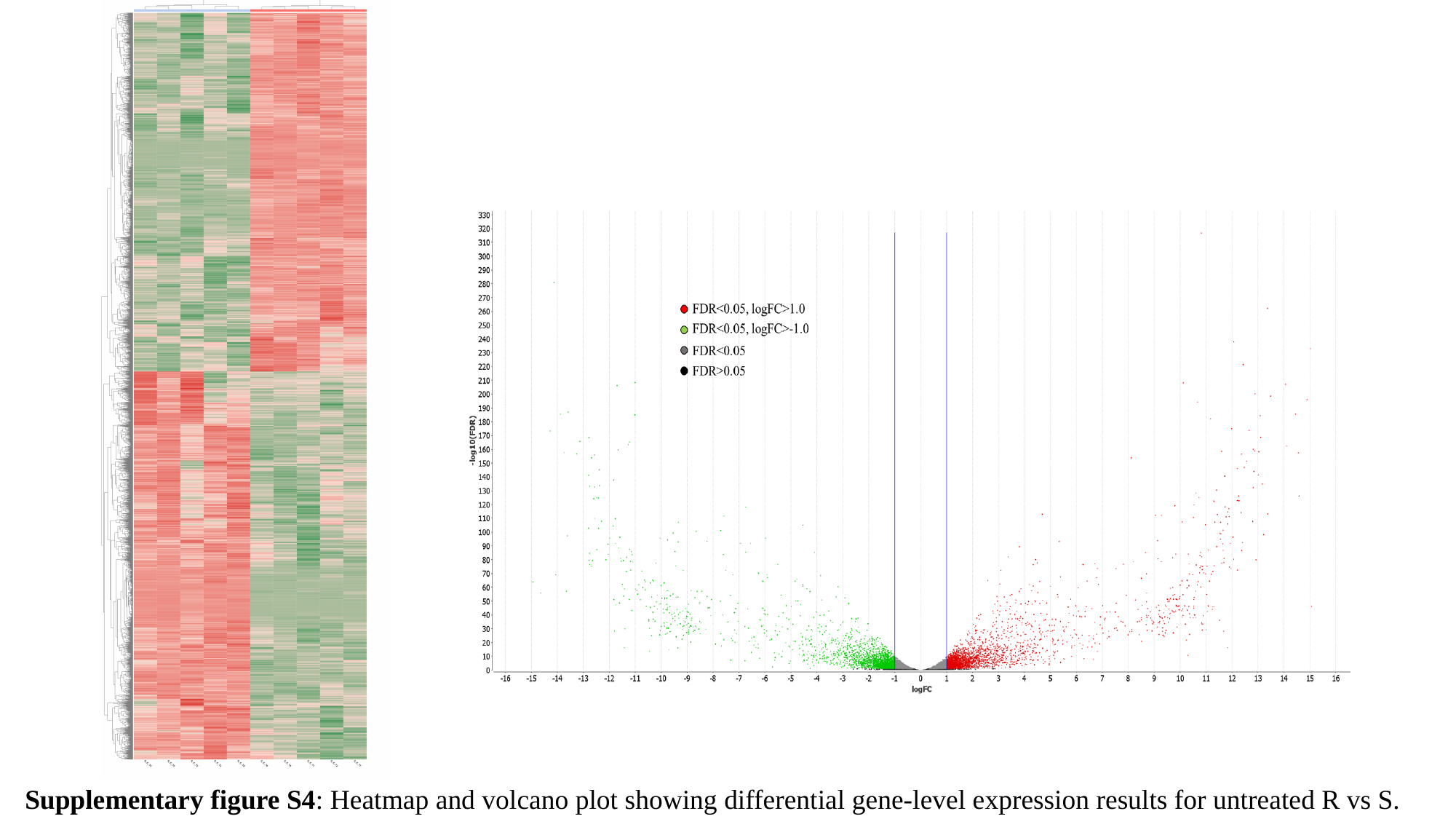

Supplementary figure S4: Heatmap and volcano plot showing differential gene-level expression results for untreated R vs S.

### Slide 6
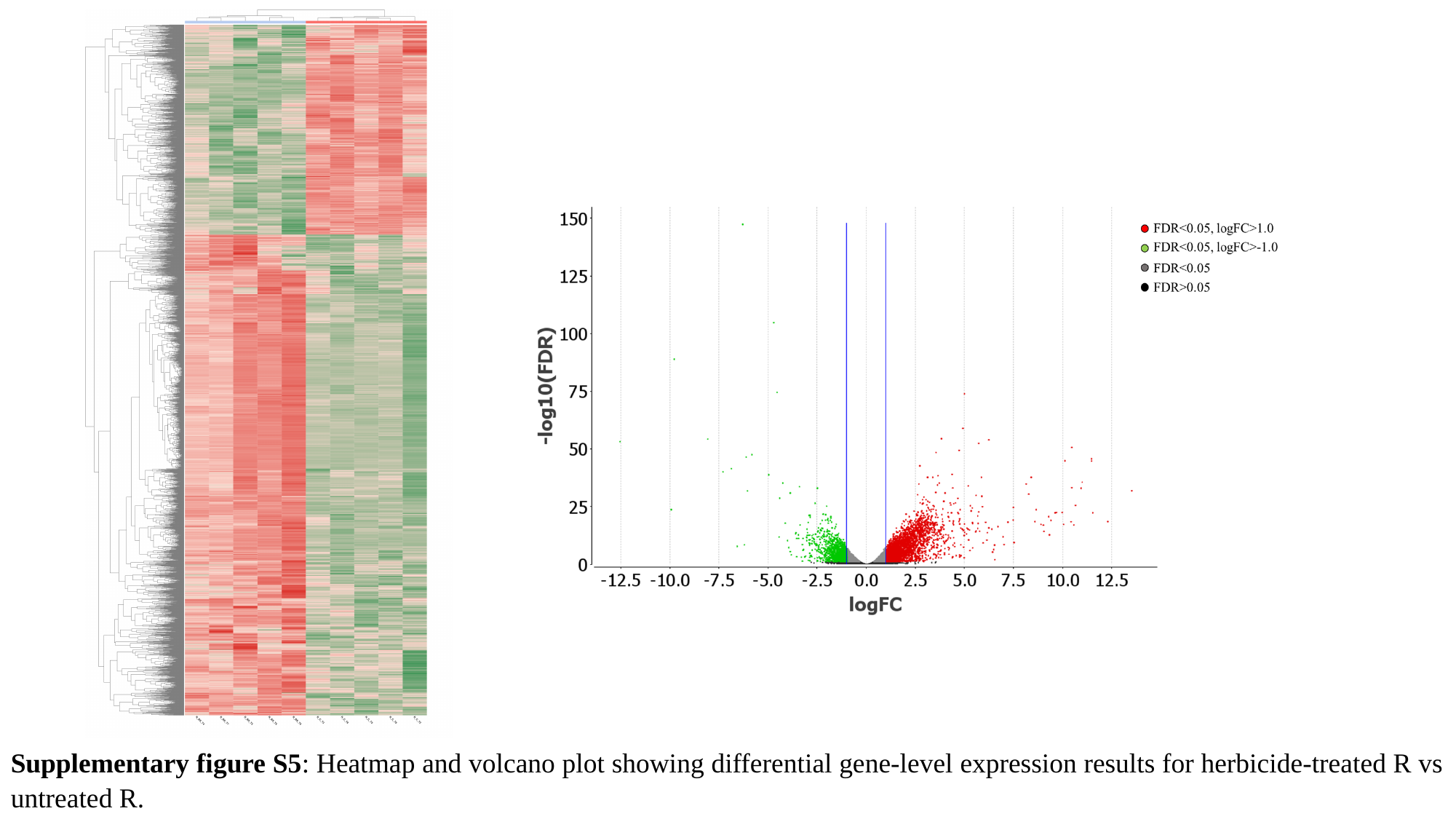

Supplementary figure S5: Heatmap and volcano plot showing differential gene-level expression results for herbicide-treated R vs untreated R.

### Slide 7
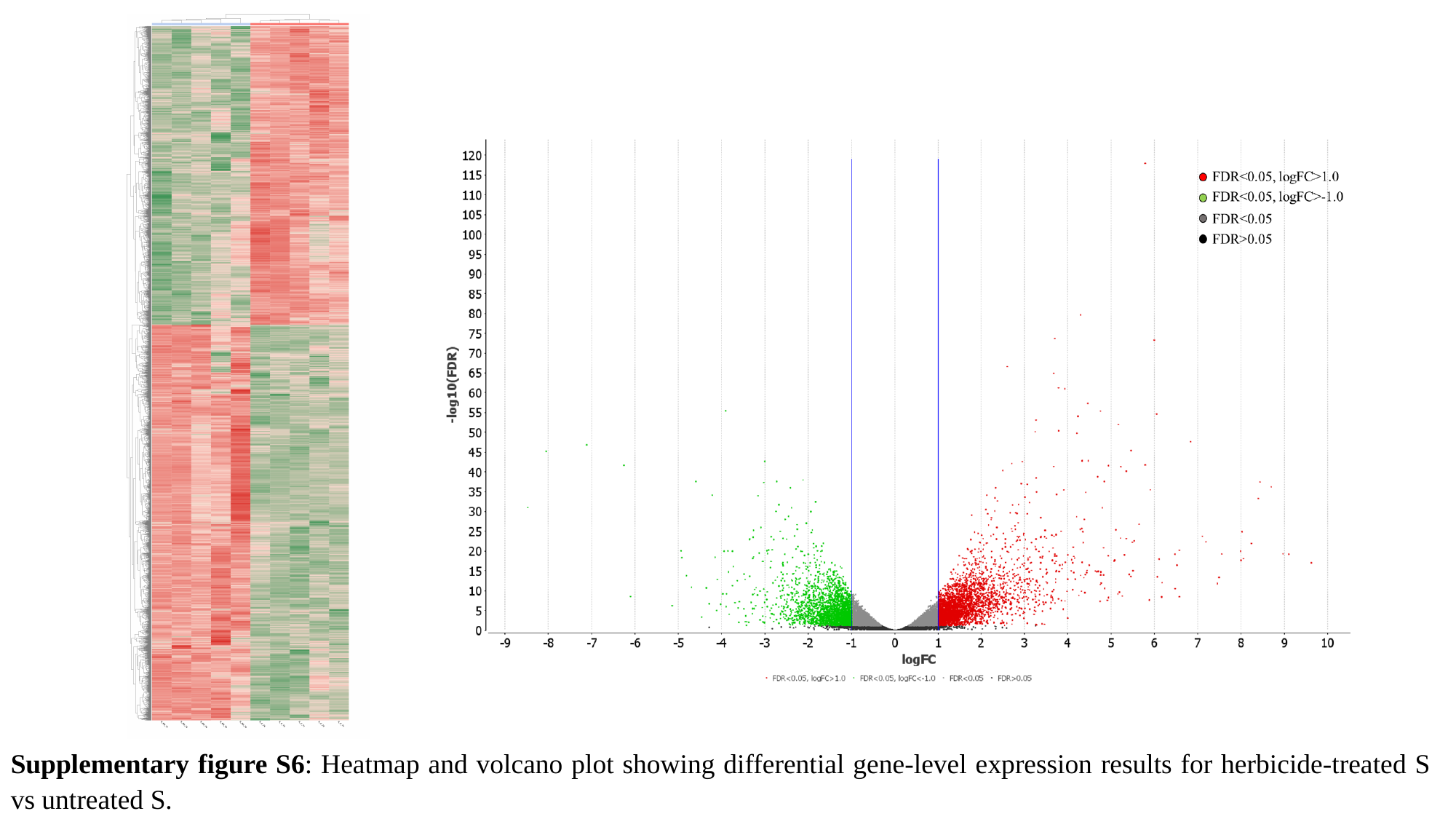

Supplementary figure S6: Heatmap and volcano plot showing differential gene-level expression results for herbicide-treated S vs untreated S.

### Slide 8
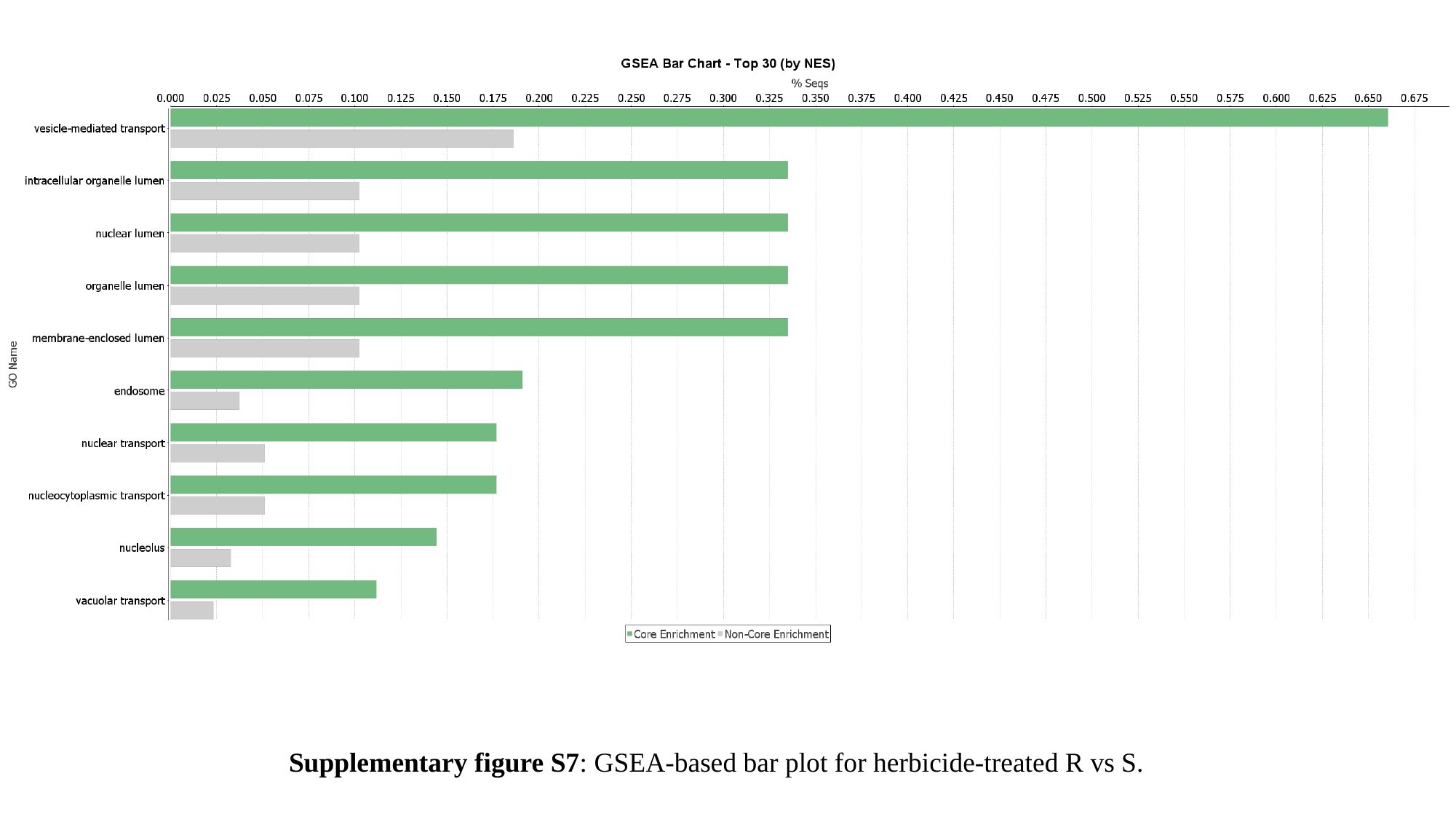

Supplementary figure S7: GSEA-based bar plot for herbicide-treated R vs S.

### Slide 9
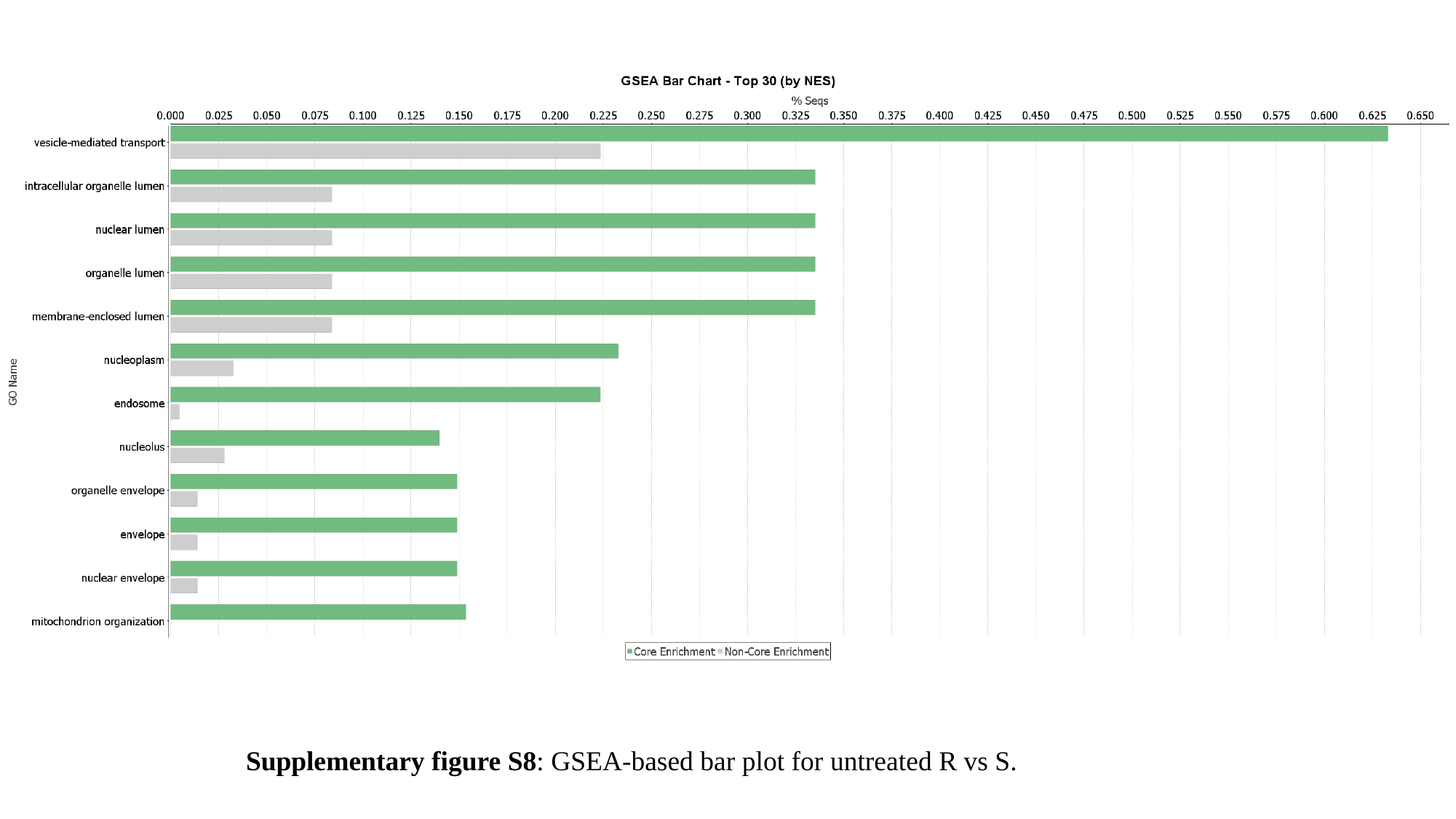

Supplementary figure S8: GSEA-based bar plot for untreated R vs S.

### Slide 10
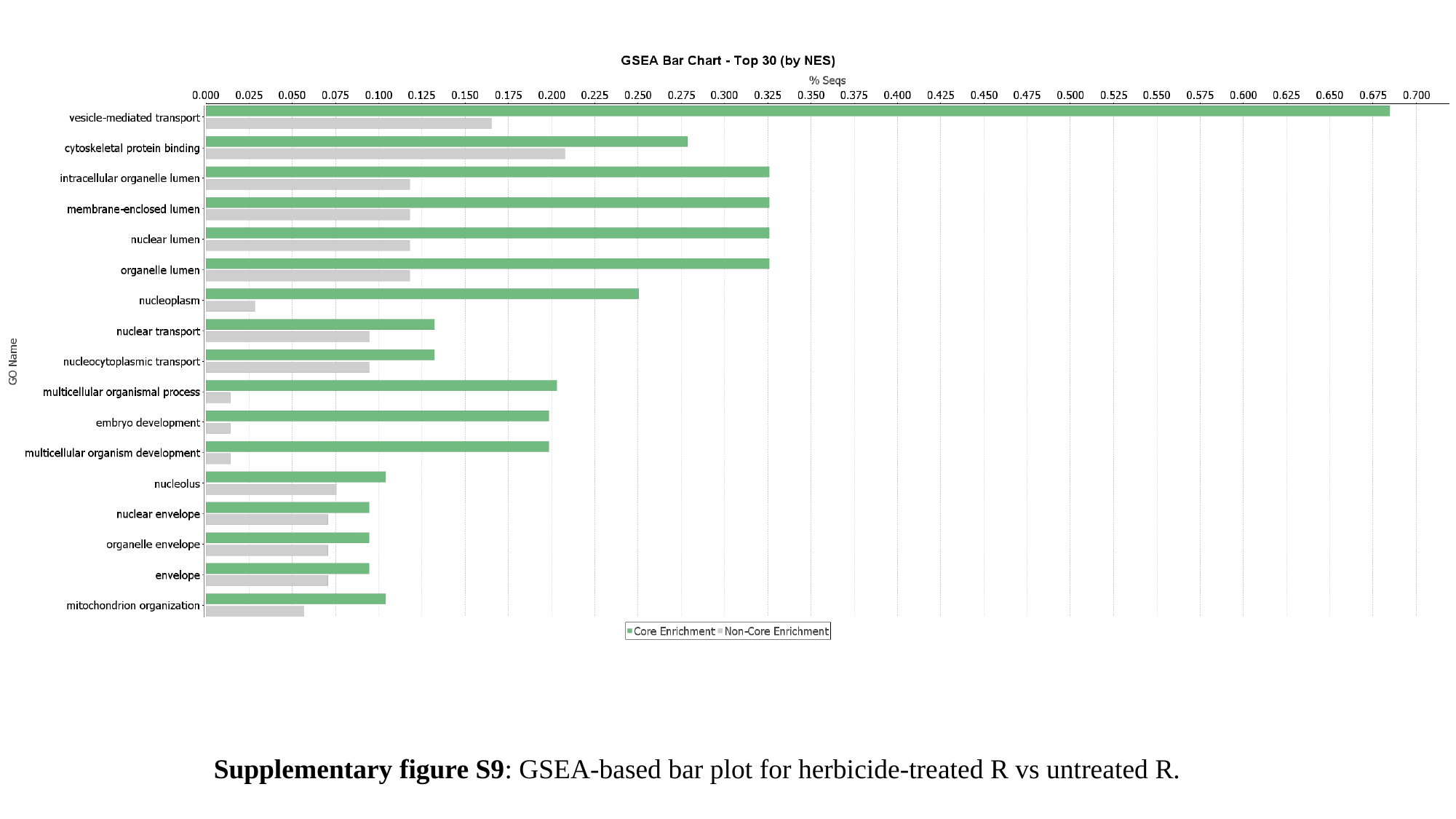

Supplementary figure S9: GSEA-based bar plot for herbicide-treated R vs untreated R.

### Slide 11
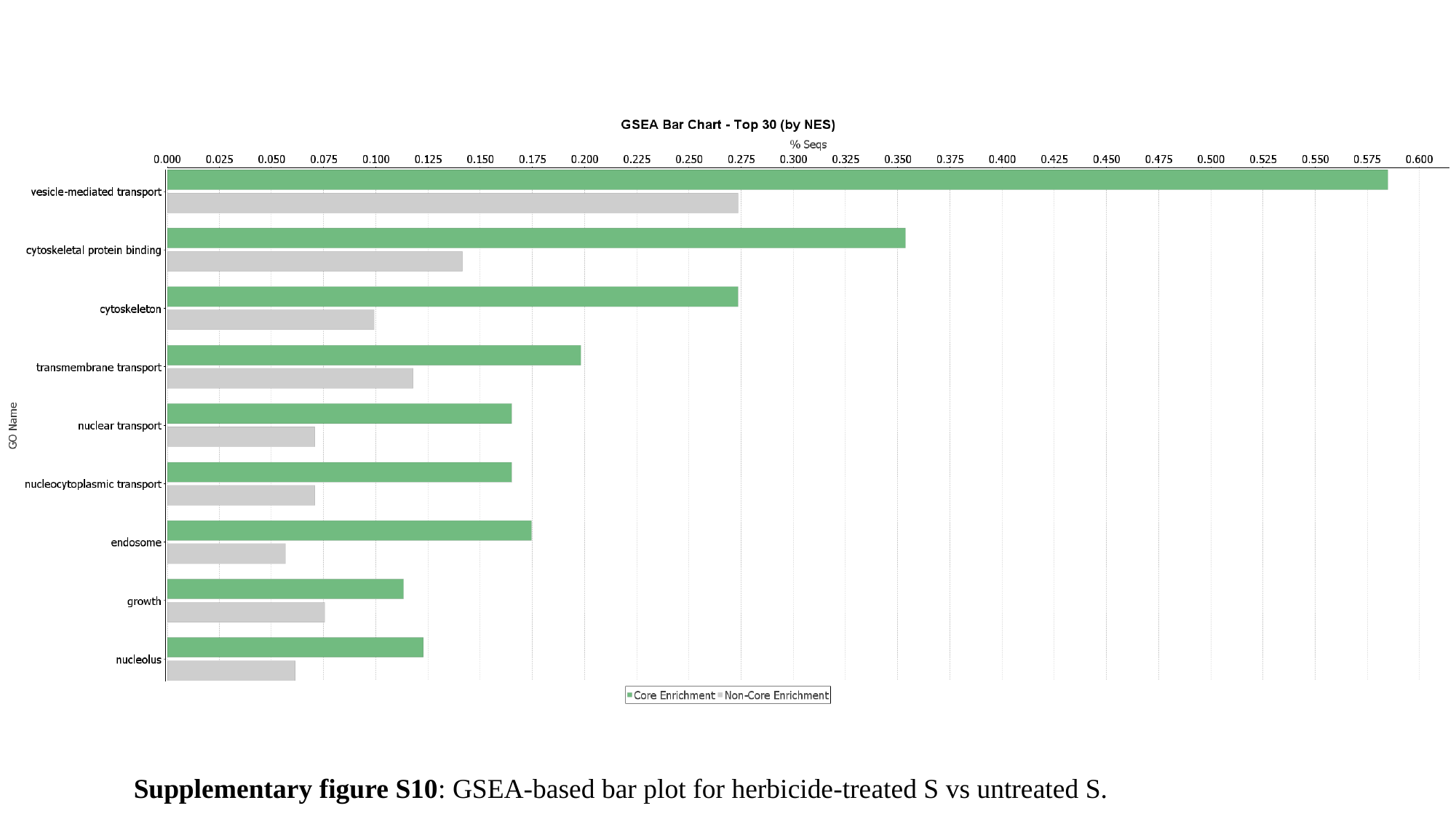

Supplementary figure S10: GSEA-based bar plot for herbicide-treated S vs untreated S.
